# An endocrine-hepato-muscular metabolic cycle links skeletal muscle atrophy and hyperglycemia in type 2 diabetes

**DOI:** 10.1101/2020.05.25.115709

**Authors:** Jürgen G. Okun, Patricia M. Rusu, Andrea Y. Chan, Yann W. Yap, Thomas Sharkie, Jonas Schumacher, Kathrin V. Schmidt, Annika Zota, Susanne Hille, Andreas Jungmann, Ludovico Maggi, Young Lee, Matthias Blüher, Stephan Herzig, Mathias Heikenwalder, Oliver Müller, Adam J. Rose

## Abstract

Both obesity and sarcopenia are frequently associated in ageing, and together may promote the progression of related conditions such as diabetes and frailty. However, little is known about the pathophysiological mechanisms underpinning this association. Here we uncover dysregulated systemic alanine metabolism and hyper-expression of the alanine transaminases (ALT) in the liver of obese/diabetic mice and humans. Hepatocyte-selective silencing of both ALT enzymes revealed a clear role in systemic alanine clearance which related to glycemic control. In obese/diabetic mice, not only did silencing both ALT enzymes retard hyperglycemia, but also reversed skeletal muscle atrophy. This was due to a rescue of depressed skeletal muscle protein synthesis, with a liver-skeletal muscle amino acid metabolic crosstalk exemplified by ex vivo experiments. Mechanistically, chronic liver glucocorticoid and glucagon signaling driven liver alanine catabolism promoted hyperglycemia and skeletal muscle wasting. Taken together, here we reveal an endocrine-hepato-muscular metabolic cycle linking hyperglycemia and skeletal muscle atrophy in type 2 diabetes.

## Introduction

Along with metabolic dysfunction and related complications, an often overlooked co-morbidity of obesity is skeletal muscle atrophy ^1,2^, which causes frailty, and is related to reduced life-quality and all-cause mortality ^3^. Nevertheless, the molecular physiology linking these two phenomena is currently poorly understood ^4^.

A hallmark of obesity-related metabolic dysfunction is hyperglycemia which is actually related to sarcopenia ^5^. Along with altered pancreatic function as well as peripheral glucose metabolism, excess hepatic glucose production is a major contributor to the hyperglycemic state in type 2 diabetes (T2D) ^6^, and strategies aimed at curtailing hepatic glucose production are effective treatment options for T2D patients ^7^. Concerning liver gluconeogenesis, several humoral substrates serve as precursors such as glycerol, pyruvate, lactate and glucogenic amino acids, with lactate and glucogenic amino acids thought to be the main contributors ^8^. In mammals, the predominant amino acid catabolized by the liver is alanine ^9,10^, and of the contribution of all 15 glucogenic amino acids to gluconeogenesis, alanine predominates ^9,11^. Furthermore, in order to clear the N-metabolites generated from catabolism of amino acids, peripheral tissues such as skeletal muscle produce alanine and glutamine as blood amino-N carriers to be subsequently taken up by the liver and gut and safely disposed of by ureagenesis, with glucose production from alanine as a result ^9,12^. This process is classically known as the glucose-alanine cycle.

This glucose-alanine cycle is dysregulated in dysglycaemic obese and T2D humans as exemplified by heightened splanchnic (i.e. viscera & liver) alanine uptake ^13,14^. These findings, generated approximately five decades ago, suggest that hepatic alanine metabolism may be an important contributor to the pathogenic hyperglycemia of T2D. Despite this, there are currently no studies which have specifically addressed the functional role of altered liver alanine metabolism in obesity and T2D. Here we demonstrate that the expression/activity of liver alanine catabolic enzymes is elevated in mouse and human T2D, the abrogation of which retards both hyperglycemia and skeletal muscle atrophy in T2D mice, and link this to endocrine regulation by the glucocorticoid and glucagon hormones.

## Results

### Heightened alanine metabolism and liver alanine aminotransferase expression/activity in obesity and type 2 diabetes

To gain further insight into whether and how amino acid metabolism is altered in obesity and type 2 diabetes (T2D), we performed blood serum amino acid profiling of two distinct mouse models of obesity-associated T2D, the BKS-db/db and New Zealand Obese mouse. In particular, we conducted this profiling from samples taken during fasting and refeeding state, as such metabolic challenges are known to reveal differences in metabolic control ^15^. While there were some changes in amino acids (Table S1), the changes in serum alanine were striking, particularly in the fasted and refed states (Fig. S1A-B). To examine this more specifically, we conducted alanine tolerance testing in lean C57Bl/6J (Bl6J; wt) and obese/diabetic BKS-db/db mice, and demonstrated that while the alanine excursion was substantially blunted (Fig. 1A), there was an exaggerated blood glucose response in the db/db mice (Fig. 1B) This suggested that alanine metabolism may be causally related to hyperglycemia in this mouse model of T2D. This prompted us to explore the molecular mechanisms behind the enhanced alanine clearance. Since the liver is known to be a key tissue that can quantitatively contribute to alanine clearance ^10^, we conducted a focused screen for transcripts known to contribute to alanine metabolism. As such, while there were only mild differences in transcripts encoding alanine transporters and alanine-glyoxylate transaminase isoforms, the alanine metabolic enzymes glutamic-pyruvic transaminase (Gpt) isoforms, otherwise more commonly known as alanine aminotransferase (ALT), were substantially higher in T2D mice (Fig. 1C). This result was confirmed at the protein level as determined by immunoblot (Fig. 1D) and immunohistochemistry (Fig. 1E), as well as liver ALT activity (Fig. 1F), in various mouse models of obesity, insulin resistance/pre-diabetes as well as frank T2D. Importantly, obese humans with T2D also have higher levels of GPT isoform transcripts when compared with obese normoglycemic counterparts (Fig. 1G; Table S2). In this cohort, liver *GPT* and *GPT2* expression correlated with biomarkers of poor metabolic health such as waist to hip ratio and visceral fatness, indices of impaired glucose homeostasis such as HbA1c, fasting plasma glucose, HOMA-IR, and glucose clamp glucose infusion rate, as well as markers of dyslipidaemia including LDL-Cholesterol and plasma triglyceride levels (Table S3). Taken together, this prompted us to investigate a functional role of GPT and GPT2 in obesity-related metabolic dysfunction.

**Figure 1.**
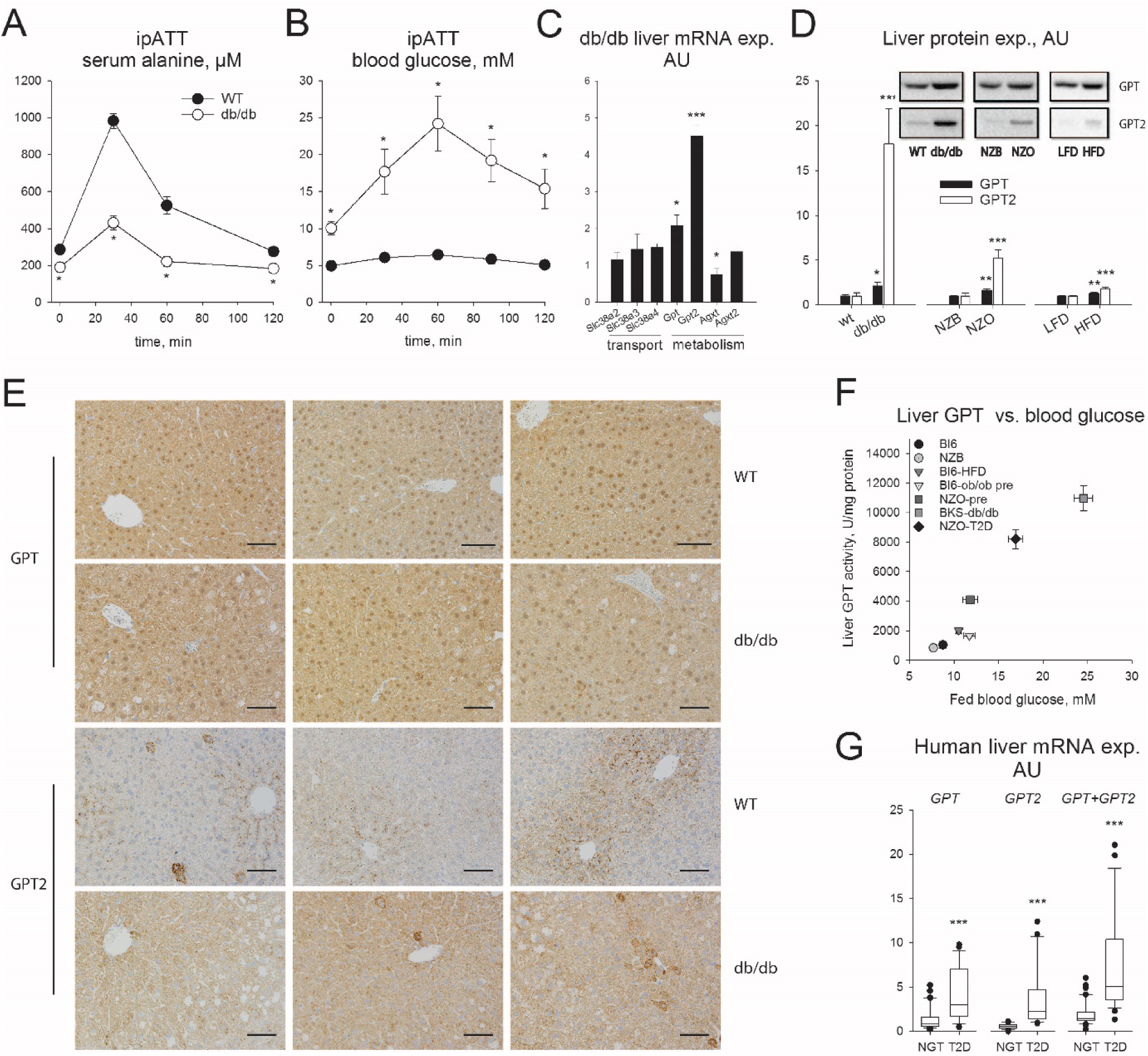
Alanine metabolism and liver alanine aminotransferase isoform expression/activity is upregulated in obesity and type 2 diabetes (T2D). A: Serum alanine levels during an intraperitoneal alanine tolerance test (ipATT) in C57Bl/6J (WT) and BKS-db/db (db/db) mice. N=4/group. Genotype difference: *P < 0.05. B: blood glucose levels from mice as in A. C: Liver messenger RNA (mRNA) expression of selected gene transcripts including solute carrier 38 (Slc38) as well as glutamic-pyruvic transaminase (Gpt) and alanine-glyoxlate aminotransferase (Agxt) isoforms in BKS-db/db vs WT mice. Data are fold expression difference of WT (C57Bl/6J). N=4/group. * genotype difference: P < 0.05. D: Liver protein expression of glutamic-pyruvic transaminase (GPT) isoforms in C57Bl/6J (WT) and BKS-db/db (db/db), New Zealand Black (NZB) and New Zealand Obese (NZO), as well as low-fat (LFD) and high-fat diet (HFD) fed C57Bl/6N mice. Data are fold expression difference of the control group. WT vs. db/db and NZB vs. NZO: N=4/group. LFD vs. HFD: n = 7-8/group. Difference versus control group: *P < 0.05, **P<0.01, ***P<0.001. E: Liver glutamic-pyruvic transaminase (GPT) immunohistochemical staining of sections taken from C57Bl/6J (WT) and BKS-db/db (db/db) mice. Shown are 3 representative images taken from 3 individual mice per group. Scale bar: 50µm. F: Liver GPT activity in various mouse models of obesity-induced insulin-resistance and T2D. Bl6: C57Bl/6J on LFD (N=8). Bl6-HFD: C57Bl/6J on HFD (N=7). NZB: New Zealand Black (N=4). NZO-pre: young pre-diabetic New Zealand Obese (N=4). NZO-T2D: older New Zealand Obese with T2D (N=6). Bl6-ob/ob-pre: young pre-diabetic obese ob/ob mice on the C57Bl/6J strain (N=5). BKS-db/db: obese-diabetic db/db mice on the BKS strain (N=4). G: Liver mRNA expression of glutamic-pyruvic transaminase (GPT) isoforms in humans with normal glucose tolerance (NGT, N=) or with T2D (N=). Difference between groups: ***P < 0.001. Statistical tests: A, B: 2-way repeated measures ANOVA with Holm-Sidak posthoc hoc tests. C, D: Students t-tests. G: Mann-Whitney U rank sum tests.

### Silencing of both, but not individual, liver alanine aminotransferase isoforms affect systemic alanine and glucose homeostasis upon diverse nutritional challenges

To initially assess the contribution of these liver enzymes to normal metabolic physiology, we cloned, screened, subcloned, and subsequently produced adeno-associated viruses (AAV) which transduce and express micro-RNAs (miR) to silence Gpt and Gpt2 transcript expression/activity in a hepatocyte-selective fashion ^16^. Importantly, the Gpt and Gpt2-specific AAV-miR silenced the transcripts in a potent and selective manner in the liver at the mRNA (Fig. S2A) and protein (Fig. S2B) level, 8 weeks following AAV-administration. No differences were found between controls and hepatocyte Gpt/Gpt2 knockdown (KD) mice in biometrics such as body mass as well as liver, muscle and adipose tissue mass (Fig. S2C-F). Despite this, somewhat surprisingly, the levels of alanine during an alanine challenge were not different in the single Gpt isoform knockdown, but were substantially higher after alanine administration when both liver transcripts were silenced (Fig. 2A). This also corresponded to a much lower increase in blood glucose upon alanine administration with the double GPT isoform knockdown (Fig. 2B). Since liver GPT expression is higher in obesity/diabetes, we then wanted to assess if liver overexpression of GPT isoforms would be sufficient to affect systemic alanine and glucose homeostasis. To this end, we mutated Ser126 and Ser153 to Arg in human GPT and GPT2 respectively, as these mutants reportedly decrease enzymatic function without affecting expression ^17^, and we cloned the Flag- and myc-tagged mutated and wildtype cDNAs into AAV vectors. Again, we could successfully silence both GPT and GPT2 in mouse livers in vivo, and could overexpress the human enzymes at the mRNA level (Fig. 2C; Fig. S2H). However, while the native human GPT isoforms could be successfully expressed, the mutant forms were expressed to a lesser extent at the protein level, despite similar mRNA levels to the wildtype forms (Fig. 2C and S2H). Regarding functional metabolic effects, silencing of GPT/2 again produced similar results as before with reduced alanine clearance and blunted glycaemia with an alanine challenge, but overexpression of human GPT/2 resulted in opposite effects, with enhanced alanine clearance but exaggerated glycaemia in response to the alanine challenge (Fig. 2D-E). These results show that liver GPT/2 expression is both sufficient and necessary to affect systemic alanine metabolism.

**Figure 2.**
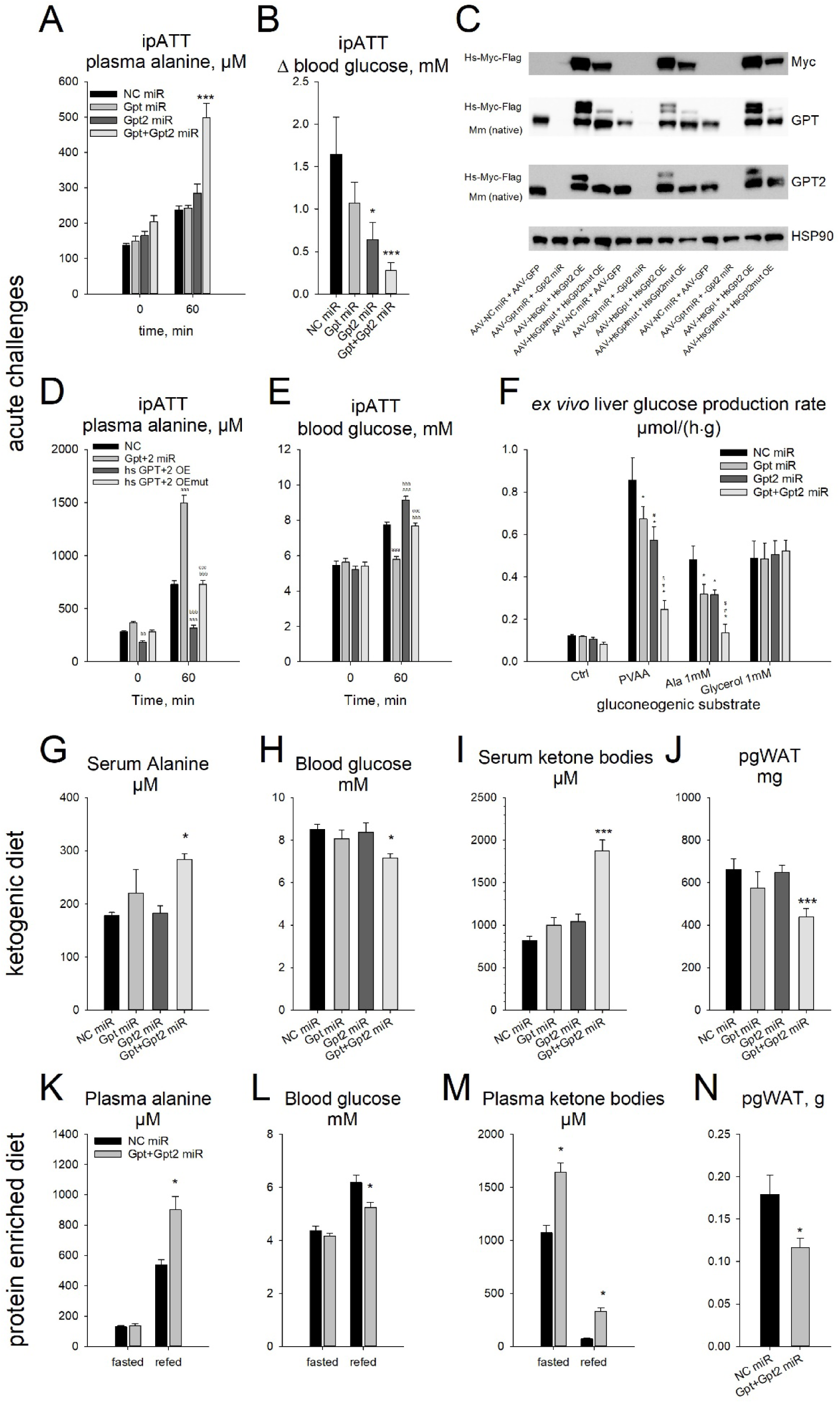
Silencing of both, but not individual, liver alanine aminotransferase isoforms affects systemic alanine and glucose homeostasis upon diverse nutritional challenges. A: Plasma alanine levels during an intraperitoneal alanine tolerance test (ipATT) in mice with hepatocyte selective AAV-miR mediated silencing of glutamic-pyruvic transaminase (Gpt) isoforms. NC: negative control. miR: micro-RNA. Data are mean ± SEM, N = 6-7/ group. Effect of miR vs. NC miR: * P < 0.05, ** P < 0.01, *** P < 0.001. B: The change () in blood glucose between time 0 and 60 from mice as in A. Data are mean ± SEM. Effect of miR vs. NC miR: * P < 0.05, ** P < 0.01, *** P < 0.001. C: Western blot of proteins Myc-tag (Myc), glutamic-pyruvic transaminase (GPT), GPT2, and the loading control heat shock protein 90 (HSP90) of liver samples from mice pre-treated with adeno-associated viruses to express a negative control micro-RNA (NC-miR) and green fluorescent protein (GFP), Gpt and GPT2-specfific miRs, human Gpt and Gpt2 mRNAs (HsGpt OE, HsGpt2 OE), and mutants of human Gopt and Gpt2 mRNAs to produce enzymatically inactive proteins (HsGptmut OE, HsGPT2mut OE). Shown are blots of 3 replicates per group. D: Plasma alanine levels during an intraperitoneal alanine tolerance test (ipATT) in mice treated as in C. Data are mean ± SEM, N = 6/ group. Different than NC: ^a^ P < 0.05, ^aa^ P < 0.01, ^aaa^ P < 0.001; different than Gpt+2 miR: ^b^ P < 0.05, ^bb^ P < 0.01, ^bbb^ P < 0.001. Different than hs GPT+2 OE: ^c^ P < 0.05, ^cc^ P < 0.01, ccc P < 0.001. E: Blood glucose levels during an intraperitoneal alanine tolerance test (ipATT) in mice treated as in C. Data are mean ± SEM, N = 6/ group. Different than NC: ^a^ P < 0.05, ^aa^ P < 0.01, ^aaa^ P < 0.001; different than Gpt+2 miR: ^b^ P < 0.05, ^bb^ P < 0.01, ^bbb^ P < 0.001. Different than hs GPT+2 OE: ^c^ P < 0.05, ^cc^ P < 0.01, ccc P < 0.001. F: *Ex vivo* glucose production rates from precision cut liver slices taken from mice with hepatocyte selective AAV-miR mediated silencing of glutamic-pyruvic transaminase (Gpt) isoforms. Shown are glucose production rates from control media (Ctrl), as well as media containing hepatic portal vein amino acid concentrations (PVAA) as well as alanine (Ala), glutamine (Gln), pyruvate (Pyr) and glycerol at the indicated concentrations. Data are from 3 separate mice per AAV averaged from data from 3 slices used per treatment (n = 9 per group total). G: Serum alanine levels in mice fed a ketogenic diet with hepatocyte selective AAV-miR mediated silencing of glutamic-pyruvic transaminase (Gpt) isoforms. NC: negative control. miR: micro-RNA. Data are mean ± SEM, N = 6/group. Effect of miR vs. NC miR: * P < 0.05, ** P < 0.01, *** P < 0.001. H: Ad libitum fed blood glucose levels from mice as in G. I: Ad libitum fed serum ketone body levels from mice as in G. J: Perigonadal white adipose tissue (pgWAT) mass of mice as in G. K: Serum alanine levels in mice fasted or refed following adaptation to a protein-enriched (80%E) diet with hepatocyte selective AAV-miR mediated silencing of glutamic-pyruvic transaminase (Gpt) isoforms. NC: negative control. miR: micro-RNA. Data are mean ± SEM, N = 6/group. Effect of miR vs. NC miR: * P < 0.05, ** P < 0.01, *** P < 0.001. L: Blood glucose level from mice as in K. M: Plasma ketone bodies from mice as in K. N: Perigonadal white adipose tissue (pgWAT) mass at the end of the experimental period from mice as in K. Statistical tests: A, D-E, K-M: 2-way repeated measures ANOVA with Holm-Sidak posthoc hoc tests. B, F-J: 1-way ANOVA with Holm-Sidak posthoc hoc tests. N: students t-test.

Conceivably, these effects could be due to liver effects on multisystemic metabolic function. To thus specifically examine liver metabolic effects of GPT/2, we assessed liver metabolism ex vivo following GPT/2 silencing in vivo via AAV administration (Fig. 2F). While the ex vivo hepatic glucose production rate in response to alanine or a mixture of amino acids corresponding to hepatic portal vein amino acid concentrations was lowered by the single Gpt isoform silencing, this lowering was greater with the double isoform silencing (Fig. 2F). Furthermore, the effects of Gpt/2 silencing in response to glycerol were negligible (Fig. 2F), demonstrating that the liver remained metabolically functional as the glycerol reaction feeds gluconeogenesis at a distal step from the Gpt/2 reactions.

Given that there were Gpt/Gpt2 dependent and independent responses of HGP to different gluconeogenic substrates, we then tested the requirement for liver alanine metabolism under a more physiological setting such as fasting and refeeding where all substrates can interact. Again serum alanine was affected, again only in the double KD group and only during refeeding (Fig. S2I). Despite this, there was no difference in blood glucose (Fig. S2J) suggesting a potential compensation by alternate gluconeogenic substrates such as other amino acids, lactate or glycerol. Indeed, while serum lactate, glycerol and non-esterified fatty acids were not different (Fig. S2K-M), serum ketone bodies were higher (Fig. S2N), suggesting a compensatory upregulation in ketogenesis to spare glucose consumption by other tissues.

Prompted by the knowledge of the potential adaptive metabolism to compensate for liver alanine catabolic capacity, we next conducted nutritional studies to unveil the dependency of alanine as a liver gluconeogenic substrate. First we conducted a ketogenic diet study whereby mice with liver ALT loss-of-function (Fig. S2O) were fed a diet devoid of carbohydrate to force the use of gluconeogenic substrates to maintain glycaemia. Indeed, even in the ad libitum fed state, serum alanine was higher (Fig. 2G) with correspondingly lower blood glucose levels (Fig. 2H). Again, despite no differences in total body mass and liver mass (Fig. S2P-Q) as well as other metabolites (Fig S2R-U), there were higher ketone body levels (Fig. 2I) with a lower gain of fat mass during the diet experiment (Fig. 2J & S2V) in mice with liver ALT silencing. In a different approach, we also employed a high-protein/low carbohydrate diet, where again the mice would be forced to increase the liver amino acid catabolic rate and ureagenesis and contribution to gluconeogenesis ^10^. During acute fasting-refeeding experiments, serum alanine was clearly higher in the refed state with liver ALT silencing (Fig. 2K), which corresponded to lower glucose in the refed (Fig. 2L) and ad libitum fed (Fig. S2W) state, with again higher ketone body levels (Fig. 2M). Again, despite no differences in body, liver and skeletal muscle mass (Fig. S2X-Z), perigonadal fat mass was lower (Fig. 2N), highlighting a compensatory upregulation of triglyceride mobilization perhaps to provide the alternate gluconeogenic substrate glycerol, as well as fatty acids for ketogenesis, to the liver. Taken together, liver alanine catabolism via ALT contributes to systemic alanine homeostasis and glycemic control in vivo, but glycemia can be maintained by either by recruitment of alternate gluconeogenic substrates and/or by upregulated ketogenesis.

### Silencing of both, but not individual, liver alanine aminotransferase isoforms retards hyperglycemia in mouse models of obesity-related type 2 diabetes

As ALT expression and activity was potently upregulated in the obese/diabetic BKS-db/db mouse (Fig. 1), we next assessed whether silencing of liver ALT activity would retard the hyperglycemia in the mice, especially as we could show that liver ALT contributes to glycemic regulation in lean mice under certain conditions (Fig. 2). Indeed, hepatic silencing (Fig. S3A-B) of both, but not individual, ALT isoforms in BKS-db/db mice reversed the exaggerated alanine tolerance in these mice (Fig 3A), which coincided with a lowering of the glycemic response to alanine (Fig. 3B), independent of changes in body-(Fig. S3C) and liver-(Fig. S3D) mass. Strikingly markers of frank T2D such as polydipsia (Fig. 3C) and glycosuria (Fig. 3D) were also lowered by liver-specific ALT silencing. In addition, in response to a refeeding challenge, there was lower blood glucose (Fig. S3E) despite no differences in food consumption (Fig. S3F). Moreover, these differences in alanine and glucose metabolism were independent from changes in other serological parameters such as triglycerides, ketone bodies, NEFA and cholesterol (Fig. S3G-J).

**Figure 3.**
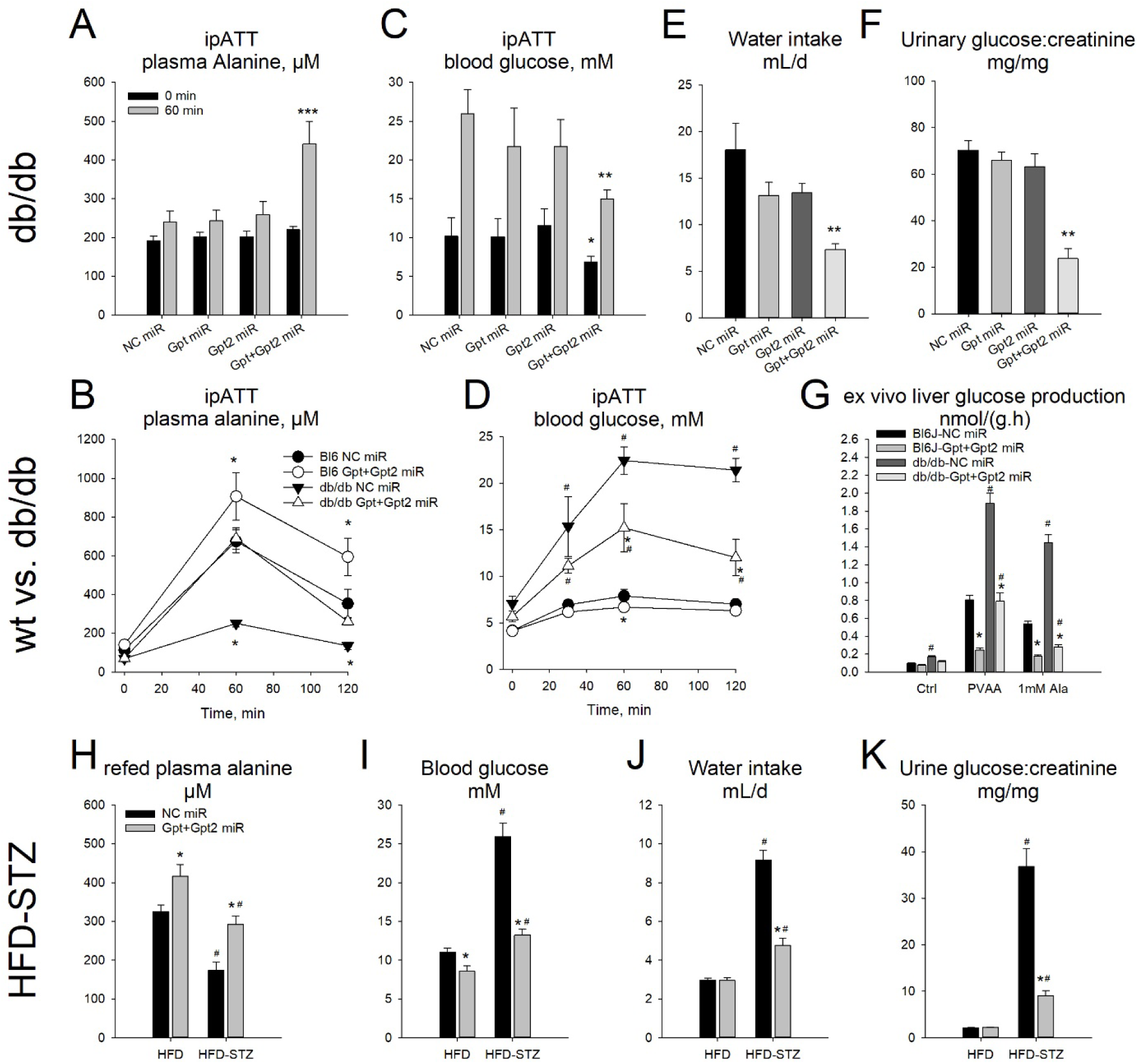
Silencing of both, but not individual, liver alanine aminotransferase isoforms retards hyperglycemia in mouse models of obesity-related type 2 diabetes. A: Plasma alanine levels during an intraperitoneal alanine tolerance test (ipATT) in obese/diabetic BKS-db/db mice with hepatocyte selective AAV-miR mediated silencing of glutamic-pyruvic transaminase (Gpt) isoforms. NC: negative control. miR: micro-RNA. Data are mean ± SEM, N = 6/group. Effect of miR vs. NC miR: * P < 0.05, ** P < 0.01, *** P < 0.001. B: Blood glucose levels of mice as in A. C: Water intake rate in mice as in A. D: Urinary glucose to creatinine ratio from mice as in A. E: Plasma alanine levels during an intraperitoneal alanine tolerance test (ipATT) in lean C57Bl/6J (Bl6) and age-matched obese/diabetic BKS-db/db mice with hepatocyte selective AAV-miR mediated silencing of glutamic-pyruvic transaminase (Gpt) isoforms. NC: negative control. miR: micro-RNA. Data are mean ± SEM, N = 4/ group. Effect of miR vs. NC miR: * P < 0.05, ** P < 0.01, *** P < 0.001. Effect of genotype: # P < 0.05, ## P < 0.01, ### P < 0.001. F: Blood glucose levels of mice as in E. G: Ex vivo glucose production rates from precision cut liver slices taken from C57Bl/6J (Bl6J) or BKS-db/db (db/db) mice with hepatocyte selective AAV-miR mediated silencing of glutamic-pyruvic transaminase (Gpt) isoforms. Shown are glucose production rates from control media (Ctrl), as well as media containing hepatic portal vein amino acid concentrations (PVAA) as well as alanine (Ala) at the indicated concentration. Data are from 3 separate mice per AAV averaged from data from 3 slices used per treatment (n = 9 per group total). H: Plasma alanine levels from 24h fasted, 6h refed mice on an obesogenic high-fat diet with (HFD-STZ) or without (HFD) streptozotocin (STZ) pre-treatment to exacerbate the progression of frank diabetes; with hepatocyte selective AAV-miR mediated silencing of glutamic-pyruvic transaminase (Gpt) isoforms. NC: negative control. Data are mean ± SEM, N = 6/group. Effect of miR vs. NC miR: * P < 0.05, ** P < 0.01, *** P < 0.001. Effect of STZ: # P < 0.05, ## P < 0.01, ### P < 0.001. I: Ad libitum fed blood glucose levels from mice in H. J: Water intake rate from mice as in H. K: Urinary glucose to creatinine ratio from mice as in H. Statistical tests: A, C, G, H, I, J, K: 2-way ANOVA with Holm-Sidak posthoc hoc tests. B, D: 2-way repeated measures ANOVA with Holm-Sidak posthoc hoc tests. E, F: 1-way ANOVA with Holm-Sidak posthoc hoc tests.

Next, we repeated these studies in C57Bl/6J and BKS-db/db mice to specifically examine the level by which we could reverse T2D compared with lean, normoglycaemic mice, and could show that liver Gpt/2 silencing (Fig. S2K-M) could revert the enhanced alanine clearance back to levels of control mice (Fig. 3E). However, again there was only a partial reversion of the exaggerated glycaemia in response to the alanine challenge (Fig. 3F). In terms of liver glucose production, similar to the responses in vivo (Fig. E), the excessive hepatic glucose production ex vivo in response to hepatic portal vein amino acids, as well as alanine alone, was abrogated by liver ALT silencing (Fig. 3G). In order to investigate this in another mouse model of T2D we employed the high-fat-diet (HFD) fed mice with low-dose streptozotocin (HFD-STZ) together with liver ALT silencing (Fig. S3N). Here the refed levels of alanine were lower in HFD-STZ mice (Fig. 3H), which coincided with the higher liver ALT activity (Fig. 3SN), and again liver ALT silencing affected systemic alanine levels (Fig. 3H). Importantly, reduced liver ALT activity also reduced ad libitum fed blood glucose levels both in HFD fed mice with impaired glucose tolerance as well as the HFD-STZ mice with frank diabetes (Fig. 3I). On this markers of frank diabetes such as polydipsia and glycosuria were substantially abrogated by liver ALT silencing (Fig. 3J-K). Taken together, these pre-clinical studies demonstrate that the enhanced liver ALT activity is causally related to the hyperglycemia in diabetes.

### Disrupting the heightened liver catabolism improves skeletal muscle size and function in mouse models of type 2 diabetes

A striking observation that we made in our initial studies was that gastrocnemius complex skeletal muscle size was higher in the BKS-db/db mice with complete liver ALT silencing (Fig. 4A). In our subsequent studies, we also could demonstrate that that liver ALT silencing reversed the reduced muscle size of obese/diabetic mice for not only gastrocnemius complex muscle, but also triceps brachii and tibialis anterior muscles (Fig. 4B-C), and this related to a reversal of skeletal muscle fiber size (Fig. 4D). This corresponded to a reversal of the reduced lean mass, with no differences in body or fat mass as assessed by ECHO-MRI (Fig. S4A-F). Importantly, this reversal of muscle size also coincided with a consistent reversal of the neuromuscular function, as assessed by forelimb grip strength testing (Fig. 4E-G). Taken together, these results demonstrated that a liver-specific metabolic process can modulate skeletal muscle quality and functionality in T2D mouse models.

**Figure 4.**
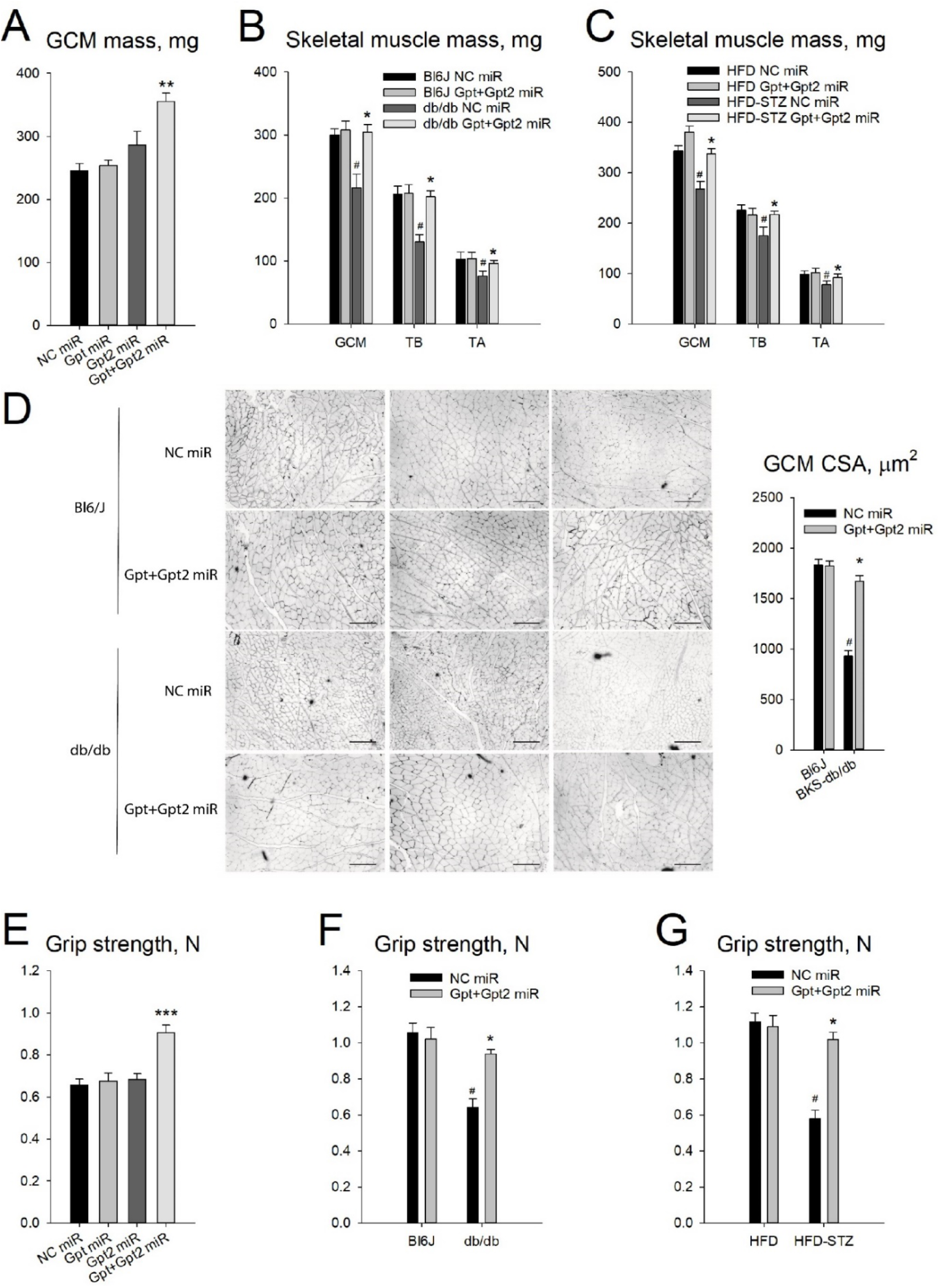
Disrupting the heightened liver alanine catabolism improves skeletal muscle size and function in mouse models of type 2 diabetes. A: Gastrocnemius complex skeletal muscle (GCM) mass in obese/diabetic BKS-db/db mice with hepatocyte selective AAV-miR mediated silencing of glutamic-pyruvic transaminase (Gpt) isoforms. NC: negative control. miR: micro-RNA. Data are mean ± SEM, N = 6/group. Effect of miR vs. NC miR: * P < 0.05, ** P < 0.01, *** P < 0.001. B: Skeletal muscle masses in lean C57Bl/6J (Bl6) and age-matched obese/diabetic BKS-db/db mice with hepatocyte selective AAV-miR mediated silencing of glutamic-pyruvic transaminase (Gpt) isoforms. NC: negative control. miR: micro-RNA. GCM: gastrocnemius complex muscle. TB: Triceps brachii. TA: tibialis anterior. Data are mean ± SEM, N = 4/group. Effect of miR vs. NC miR: * P < 0.05, ** P < 0.01, *** P < 0.001. Effect of genotype: # P < 0.05, ## P < 0.01, ### P < 0.001. C: Skeletal muscle masses in mice on an obesogenic high-fat diet with (HFD-STZ) or without (HFD) streptozotocin (STZ) pre-treatment to exacerbate the progression of frank diabetes; with hepatocyte selective AAV-miR mediated silencing of glutamic-pyruvic transaminase (Gpt) isoforms. NC: negative control. GCM: gastrocnemius complex muscle. TB: Triceps brachii. TA: tibialis anterior. Data are mean ± SEM, N = 6/group. Effect of miR vs. NC miR: * P < 0.05, ** P < 0.01, *** P < 0.001. Effect of STZ: # P < 0.05, ## P < 0.01, ### P < 0.001. D: Representative dystrophin immunohistochemical stains of fixed gastrocnemius muscle to demonstrate cross-sectional muscle fiber size from mice as in B. Shown are 3 representative images taken from 3 individual mice per group. Scale bar: 200µm. Shown to the right is the median cross sectional area (CSA). Data are mean ± SEM, N = 4/group. Effect of miR vs. NC miR: * P < 0.05, ** P < 0.01, *** P < 0.001. Effect of genotype: # P < 0.05, ## P < 0.01, ### P < 0.001. E: Forelimb grip strength from mice as in A. F: Forelimb grip strength from mice as in B. G: Forelimb grip strength from mice as in C. Statistical tests: A, E: 1-way ANOVA with Holm-Sidak posthoc hoc tests. B, C, D, F, G: 2-way ANOVA with Holm-Sidak posthoc hoc tests.

### A glucocorticoid/liver glucocorticoid receptor axis links heightened liver alanine catabolism with hyperglycemia and muscle atrophy in type 2 diabetes

Since it is known that glucocorticoids can affect both glycaemia and skeletal muscle atrophy ^18^, we investigated this as a potential mechanism behind the enhanced liver ALT driven systemic effects in obesity/diabetes. Indeed, serum corticosterone levels were higher in the two preclinical mouse models of T2D employed (Fig. 5A-B), but not obese/diabetic NZO mice (Fig. S5A), which are larger and do not exhibit muscle atrophy ^19^. In our human cohort, liver *GPT* and *GPT2* expression correlated with serum cortisol (Table S3). Further in line with a role for high glucocorticoid hormones, liver ALT isoform expression (Fig. 5C) and activity (Fig. S5B) were higher in C57Bl/6J mice chronically treated with the synthetic glucocorticoid dexamethasone (Dex), a selective activator of the glucocorticoid receptor (GR; ^18^). Hence, we conducted a study to examine whether the heightened liver ALT activity is related to the systemic effects of chronic glucocorticoid levels. As such, we could demonstrate that the higher blood glucose (Fig. S5D) as well as lower skeletal muscle size (Fig. S5E) and strength (Fig. S5F) were abrogated with liver ALT isoform silencing (Fig. S5C) upon chronic Dex treatment. This was paralleled by an abrogation in the lower body mass, with a lower liver mass and no difference in adipose tissue mass with liver ALT silencing upon chronic Dex treatment (Fig. S5G-I).

**Figure 5:**
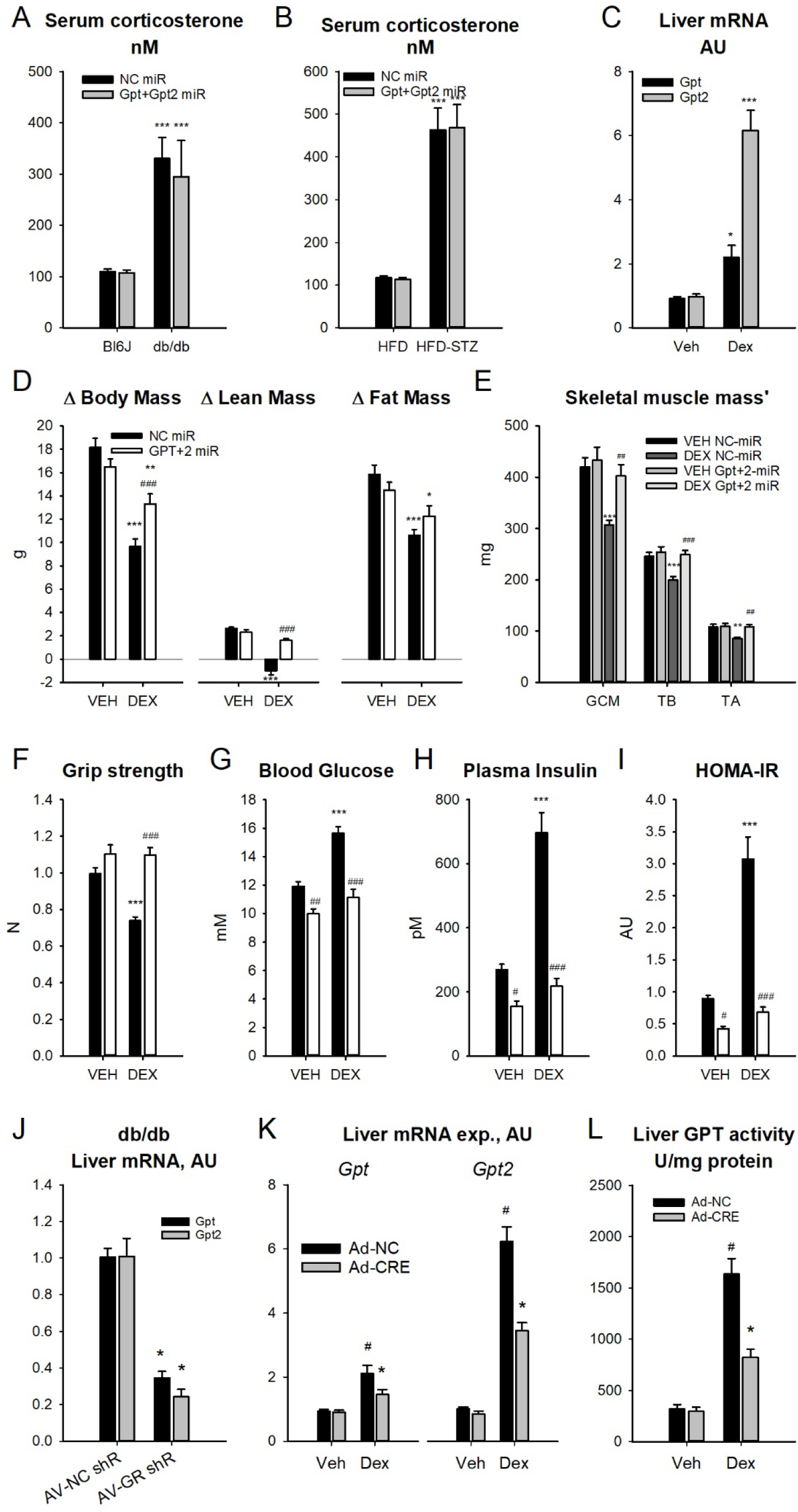
A glucocorticoid-liver glucocorticoid receptor axis links heightened liver alanine catabolism with hyperglycemia and muscle atrophy in type 2 diabetes. A: Serum corticosterone levels in lean C57Bl/6J (Bl6) and age-matched obese/diabetic BKS-db/db mice with hepatocyte selective AAV-miR mediated silencing of glutamic-pyruvic transaminase (Gpt) isoforms. NC: negative control. miR: micro-RNA. Data are mean ± SEM, N = 4/group. Effect of genotype: * P < 0.05, ** P < 0.01, *** P < 0.001. Effect of miR vs. NC miR: # P < 0.05, ## P < 0.01, ### P < 0.001. B: Serum corticosterone levels in mice on an obesogenic high-fat diet with (HFD-STZ) or without (HFD) streptozotocin (STZ) pre-treatment to exacerbate the progression of frank diabetes; with hepatocyte selective AAV-miR mediated silencing of glutamic-pyruvic transaminase (Gpt) isoforms. NC: negative control. Data are mean ± SEM, N = 6/group. Effect of STZ: * P < 0.05, ** P < 0.01, *** P < 0.001. Effect of miR vs. NC miR: # P < 0.05, ## P < 0.01, ### P < 0.001. C: Liver glutamic-pyruvic transaminase (Gpt) isoform mRNA expression in mice chronically treated with the synthetic glucocorticoid dexamethasone (Dex; 1mg/kg per day, 14d) or vehicle control (Veh). Data are mean ± SEM, N = 7/group. Effect of Dex: * P < 0.05, ** P < 0.01, *** P < 0.001. D: The change (Δ) in body mass, lean mass and fat mass in mice co-treated with a high-fat diet with or without the synthetic glucocorticoid dexamethasone (Dex; 1mg/kg per day, 14d) or vehicle control (Veh) for 6 weeks, with hepatocyte selective AAV-miR mediated silencing of glutamic-pyruvic transaminase (Gpt) isoforms. Data are mean ± SEM, N = 8/group. Effect of Dex: # P < 0.05, ## P < 0.01, ### P < 0.001. # P < 0.05, ## P < 0.01, ### P < 0.001. E: Mass’ of skeletal muscles including gastrocnemius complex (GCM), triceps brachii (TB), and tibialis anterior (TA) of mice as in D. F: Forelimb grip strength of mice as in D. G: Blood glucose of mice as in D. H: Plasma insulin levels of mice as in D. I: Homeostatic model assessment of insulin resistance (HOMA-IR) of mice as in D. J: Liver Gpt isoform mRNA expression of BKS-db/db mice with liver-specific silencing of the glucocorticoid receptor (GR) via adenoviral mediated transduction an expression of a specific shRNA (AV-GR shR) or a negative control shRNA (AV-NC shR). Data are mean ± SEM, N = 4/group. Effect of GR shR vs. NC shR: * P < 0.05, ** P < 0.01, *** P < 0.001. K: Liver Gpt isoform mRNA expression of GR-floxed mice pre-treated with adenoviral constructs expressing (Ad-CRE) or not (Ad-NC) Cre-recombinase with (Dex) or without (Veh) chronic dexamethasone treatment. Data are mean ± SEM, N = 6–7/group. Effect of Ad-CRE vs. Ad-NC: * P < 0.05, ** P < 0.01, *** P < 0.001. Effect of Dex: # P < 0.05, ## P < 0.01, ### P < 0.001. L: Liver GPT activity of mice as in K. Statistical tests: A, B, D, E, F, G, H, I, K, L: 2-way ANOVA with Holm-Sidak posthoc hoc tests. C, J: Students t-tests. D: Blood glucose of C57Bl/6J in mice chronically treated with the synthetic glucocorticoid dexamethasone (Dex; 1mg/kg per day, 14d) or vehicle control (Veh) with hepatocyte selective AAV-miR mediated silencing of glutamic-pyruvic transaminase (Gpt) isoforms. NC: negative control. miR: micro-RNA. Data are mean ± SEM, N = 8/group. Effect of miR vs. NC miR: * P < 0.05, ** P < 0.01, *** P < 0.001. Effect of Dex: # P < 0.05, ## P < 0.01, ### P < 0.001. E: Skeletal muscle masses of mice as inE. GCM: gastrocnemius complex muscle. TB: Triceps brachii. TA: tibialis anterior. F: Forelimb grip strength of mice as in E. G: Liver Gpt isoform mRNA expression of BKS-db/db mice with liver-specific silencing of the glucocorticoid receptor (GR) via adenoviral mediated transduction an expression of a specific shRNA (AV-GR shR) or a negative control shRNA (AV-NC shR). Data are mean ± SEM, N = 4/group. Effect of GR shR vs. NC shR: * P < 0.05, ** P < 0.01, *** P < 0.001. H: Liver Gpt isoform mRNA expression of GR-floxed mice pre-treated with adenoviral constructs expressing (Ad-CRE) or not (Ad-NC) Cre-recombinase with (Dex) or without (Veh) chronic dexamethasone treatment. Data are mean ± SEM, N = 6–7/group. Effect of Ad-CRE vs. Ad-NC: * P < 0.05, ** P < 0.01, *** P < 0.001. Effect of Dex: # P < 0.05, ## P < 0.01, ### P < 0.001.

Given this preliminary data, we then employed a model of combined high-fat diet feeding with chronic dexamethasone treatment which is known to produce a severe manifestation of the metabolic syndrome in rodents ^20–22^. In particular, we combined this model with hepatocyte-silencing of ALT isoforms, to determine whether this would affect susceptibility to metabolic disease and atrophy in this model. While, DEX treatment depressed the gain in total mass and lean mass, this was abrogated with liver GPT/2 silencing (Fig. 5D). In terms of tissue/organ mass, the lower skeletal muscle mass seen upon DEX treatment was completely abrogated by liver GPT/2 silencing. In addition, while there were no effects on perigonadal fat, the higher liver mass with combined HFD and DEX treatment was also abrogated by liver GPT/2 silencing (Fig SJ). Similar to prior studies using other T2D models (Fig 4), the changes in muscle mass were reflected in functionality as assessed by grip strength testing (Fig. 5F), and correlated with improved glucose homeostasis as assessed by blood glucose and insulin levels, as well as HOMA-IR (Fig. 5G-I).

Since the BKS-db/db mice have higher serum glucocorticoid levels (Fig. 5A), we tested whether the liver GR was a transcriptional control point behind the higher liver ALT in this setting, and could show that liver GR silencing abrogates the heightened liver ALT isoform expression in obesity/T2D mice (Fig. 5J). Furthermore, liver GR silencing blunted the increase in ALT isoform expression (Fig. 5K) and activity (Fig. 5L) upon chronic Dex treatment. Lastly, cistrome mining of publically available next-generation sequencing data ^23^, demonstrated that the GR binds to regions proximal to the transcriptional start site of the Gpt and Gpt2 genes and that this aligns with regions of CEBPβ binding as well as DNAse hypersensitivity (Fig. S5K). Taken together, these data suggest that the heightened liver ALT expression/activity, and subsequent systemic effects in T2D, is mediated by increased glucocorticoid levels and liver GR transcriptional activity.

### Chronic liver glucagon signaling affects hyperglycaemia and muscle atrophy via liver alanine catabolism

Interestingly, hepatic GR silencing did not completely block the increase in liver GPT/2 expression/activity by dexamethasone treatment (Fig. 5K-L), suggesting that other mechanisms can contribute. Indeed, muscle GR activity is required for chronic glucocorticoid effects on muscle atrophy ^24,25^, which results in raised blood levels of certain amino acids ^26^. Given that raised amino acids can trigger pancreatic glucagon (GCG) secretion^26–28^, and chronic glucagon treatment can reduce lean body mass ^29,30^, we sought to investigate the role of GCG in T2D atrophy.

Initially, we tested blood serum levels of GCG in our obese/T2D mouse models, and those exhibiting profound hyperglycaemia and skeletal muscle atrophy exhibited high GCG levels (Fig. 6A). Importantly, ex vivo treatment of liver slices with GCG increased both Gpt and Gpt2 mRNA expression, with combined GCG and DEX treatment resulting in an even greater increase in Gpt2 (Fig. S6A). In addition, treatment of T2D BKS-db/db mice with a glucagon receptor antagonist which reduces hyperglycaemia ^31^ blunted GPT/2 expression/ activity (Fig. 6B and S6B). To further test the role of glucagon, we then employed a glucagon peptide (i.e. acyl-GCG) with enhanced selectivity for the glucagon receptor and greater in vivo stability and half-life ^32^. Importantly, similar what is known ^33^, whereas acyl-GCG affected liver signaling potently, it did not affect skeletal muscle, owing to that the GCG receptor is not expressed in skeletal muscle (Fig. S6C). This then makes a chronic GCG challenge an excellent model to study the role of hepatic alanine catabolism in muscle atrophy, particular as GCG potently affects liver amino acid metabolism and is elevated in T2D ^34^. Thus, to initially examine the role of GPT/2 in GCG responses, we conducted an acute GCG challenge in mice and could show that GPT/2 silencing reduced, whereas GPT/2 overexpression exacerbated the glycemic response (Fig. 6C). In a subsequent chronic acyl-GCG challenge, hepatic GPT/2 silencing prevented the decrease in body mass (Fig. 6D-E), owing to blunted lowering of both lean and fat mass (Fig. 6E), with little change in liver of perigonadal white adipose tissue (Fig. S6D). Concerning lean mass, chronic acyl-GCG treatment resulted in lower skeletal muscle mass (Fig. 6F) which again was blunted by hepatic GPT/2 silencing (Fig. 6F), with a similar pattern of change in neuromuscular function as assessed by grip strength (Fig. 6G). Similar to prior studies, these changes in skeletal muscle mass coincided with improved metabolic homeostasis, as the expected ^32^ hyperglycaemia with chronic acyl-GCG was completely abrogated by hepatic GPT/2 silencing (Fig. 6H).

**Figure 6.**
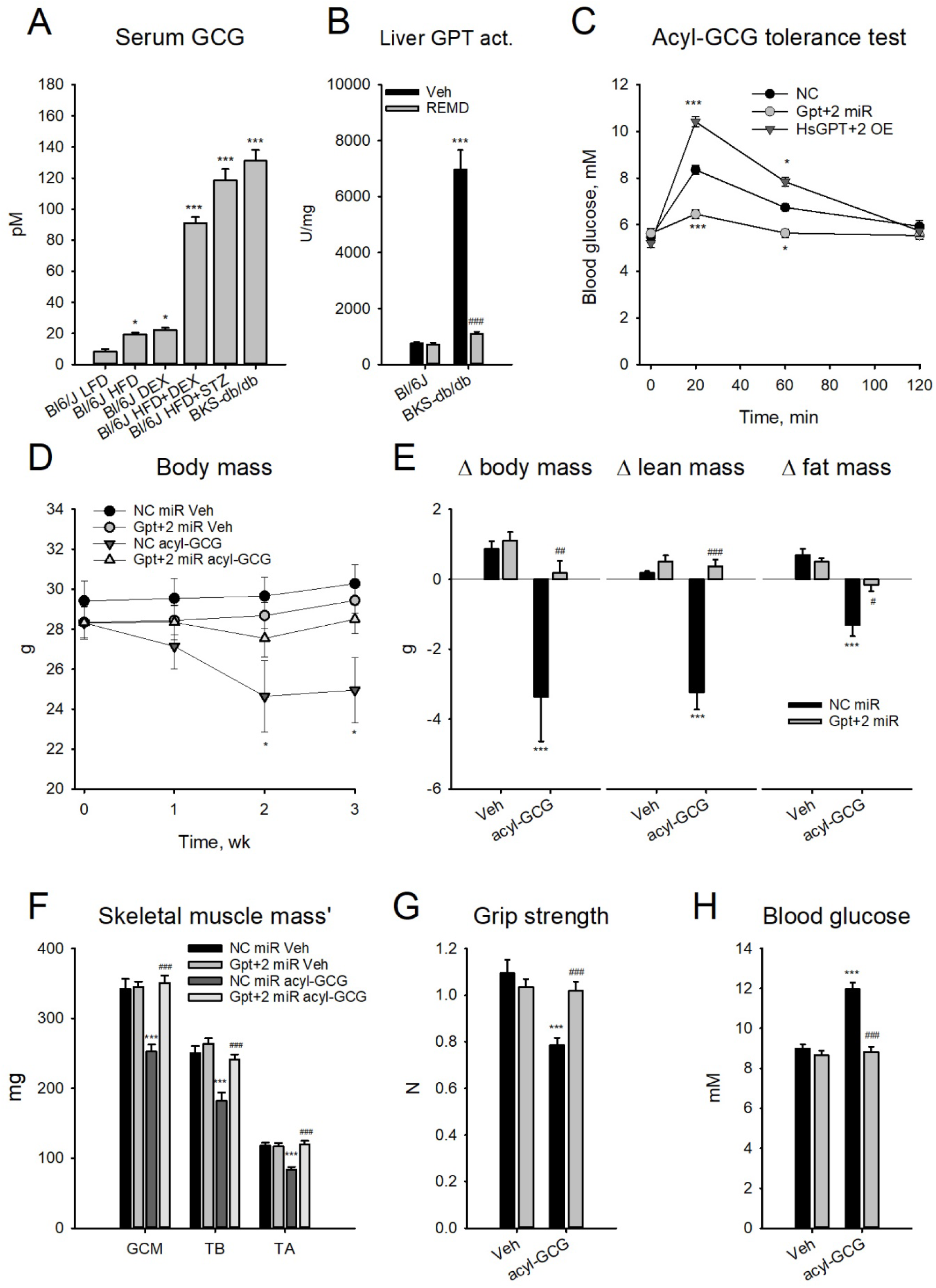
Chronic liver glucagon action affects hyperglycaemia and skeletal muscle atrophy via hepatic alanine catabolism. A: Serum glucagon (GCG) levels of mice at varying degrees of obesity and type 2 diabetes. LFD: low fat diet. HFD: high fat diet. DEX: chronic dexamethasone treatment. STZ: streptozotocin treatment. Bl6/J: C57Bl6/J mice. BKS-db/db: C57BKS mice with homozygous leptin receptor mutation. N = 4/group. B: Liver glutamic-pyruvic transaminase (GPT) activity of C57Bl6/J (Bl6/J) and obese/diabetic C57BKS mice with homozygous leptin receptor mutation (BKS-db/db) chronically treated with a glucagon receptor antagonist (REMD). N = 4/group. Different than Bl6/J LFD: * P < 0.05, ** P < 0.01, *** P < 0.001. C: Blood glucose levels of C57Bl6/J mice pre-treated with AAVs to silence hepatic Gpt isoforms (Gpt+2 miR), or overexpress human Gpt isoforms (HsGPT+2 OE) and acutely treated with acyl-glucagon (acyl-GCG). Effect of AAV vs NC (AAV NC miR + AAV GFP): * P < 0.05, ** P < 0.01, *** P < 0.001. D: Body mass’ of mice of C57Bl/6J mice chronically treated with the acyl-glucagon (acyl-GCG; 1 nmol/g/d) or vehicle control (Veh) with hepatocyte selective AAV-miR mediated silencing of glutamic-pyruvic transaminase (Gpt) isoforms. NC: negative control. miR: micro-RNA. Data are mean ± SEM, N = 8/group. Effect of miR vs. NC miR: * P < 0.05, ** P < 0.01, *** P < 0.001. Effect of Dex: # P < 0.05, ## P < 0.01, ### P < 0.001. E: The change (Δ) in body mass, lean mass and fat mass in mice as in D. F: Skeletal muscle masses of mice as in D. GCM: gastrocnemius complex muscle. TB: Triceps brachii. TA: tibialis anterior. G: Forelimb grip strength of mice as in D. H: Blood glucose levels of mice as in D. Statistical tests: A: 1-way ANOVA with Holm-Sidak posthoc hoc tests. B, E, F, G, H: -way ANOVA with Holm-Sidak posthoc hoc tests. C: 1-way repeated measures ANOVA with Holm-Sidak posthoc hoc tests. D: 2-way repeated measures ANOVA with Holm-Sidak posthoc hoc tests.

### Disrupting the heightened liver alanine catabolism reverses the reduced skeletal muscle branched chain amino acid levels and protein synthesis in type 2 diabetes

Conceivably, the lower skeletal muscle mass in diabetes could result from lower protein synthesis or greater protein breakdown ^35^. Given that known intracellular skeletal muscle proteolysis factors that are elevated in T2D ^36,37^ were not affected (Fig. S7A), we assessed *in vivo* skeletal muscle protein synthesis in normal and T2D mice. We chose to examine this only 10d after AAV administration, a time at which there was no difference in skeletal muscle mass (Fig. S7B), in order to capture a dynamic change. While skeletal muscle protein synthesis was lower in mice with obesity/diabetes, this was indeed reversed by liver ALT silencing (Fig. 7A, S7C). Importantly, there was no effect of the hepatocyte-specific AAVs on skeletal muscle ALT expression (Fig. S7D). Since it is known that there is liver-skeletal muscle metabolic crosstalk ^12^, we decided to investigate this possibility. In novel *ex vivo* co-culture experiments, we could demonstrate that this accelerated metabolic cycle was intact *ex vivo*, with liver from the obese/diabetic mice, but not Bl6J mice, depressing skeletal muscle protein synthesis rate regardless of the strain origin of the muscle (Fig. 7B), which could be reversed by liver ALT silencing (Fig. 7C). Conceivably this effect of obese/diabetic liver on skeletal muscle protein synthesis could be a result of a liver secreted factor or a metabolic interaction, so we investigated this by performing media dialysis experiments in order to normalize media metabolites but leave large molecules such as peptides in place. As such, pre-cultured and dialyzed media from obese/diabetic liver did not depress skeletal muscle protein synthesis (Fig. S7E), indicating that it is more likely to have a metabolic basis. Indeed, liver-derived factors known to influence skeletal muscle proteostasis such as insulin like growth factor 1 (Fig. S7F), follistatin (Fig. S7G), and fibroblast growth factor 21 (Fig. S7H) were largely unchanged, with lower levels of insulin (Fig. S7I), which is known to be a muscle anabolic factor ^35^. As such, we hypothesized that the heightened liver catabolism of alanine would create a ‘pull’ mechanism on muscle alanine thereby accelerating the partial catabolism of precursor amino acids in muscle for alanine synthesis ^38^. In support of this idea, of the amino acids measured, the branched-chain amino acids (BCAA), which are known to contribute quantitatively to muscle alanine output ^38^, were substantially lower in skeletal muscle of obese/diabetic mice and this was reversed by liver ALT silencing (Fig. 7D-E, S7J), and feeding excess alanine to liver-muscle co-incubates *ex vivo* reversed the effect of T2D liver on the depression of muscle protein synthesis (Fig. S7K). Lastly, to test the involvement of enhanced muscle branched chain amino acid catabolism we took advantage of the fact that muscle, but not liver hepatocytes, expresses branched chain amino acid transaminase 2 (BCAT2; neither express BCAT1) ^39^ and a novel BCAT2 inhibitor, Co8b ^40^. As such, treatment of muscle-liver co-incubates with Co8b reversed the effect of T2D liver on the depression of muscle protein synthesis (Fig. 7F). In summary, our results demonstrate that skeletal muscle protein synthesis is depressed in obesity/diabetes due to an increased pull of BCAA transamination products (e.g. alanine) from muscle towards liver.

**Figure 7.**
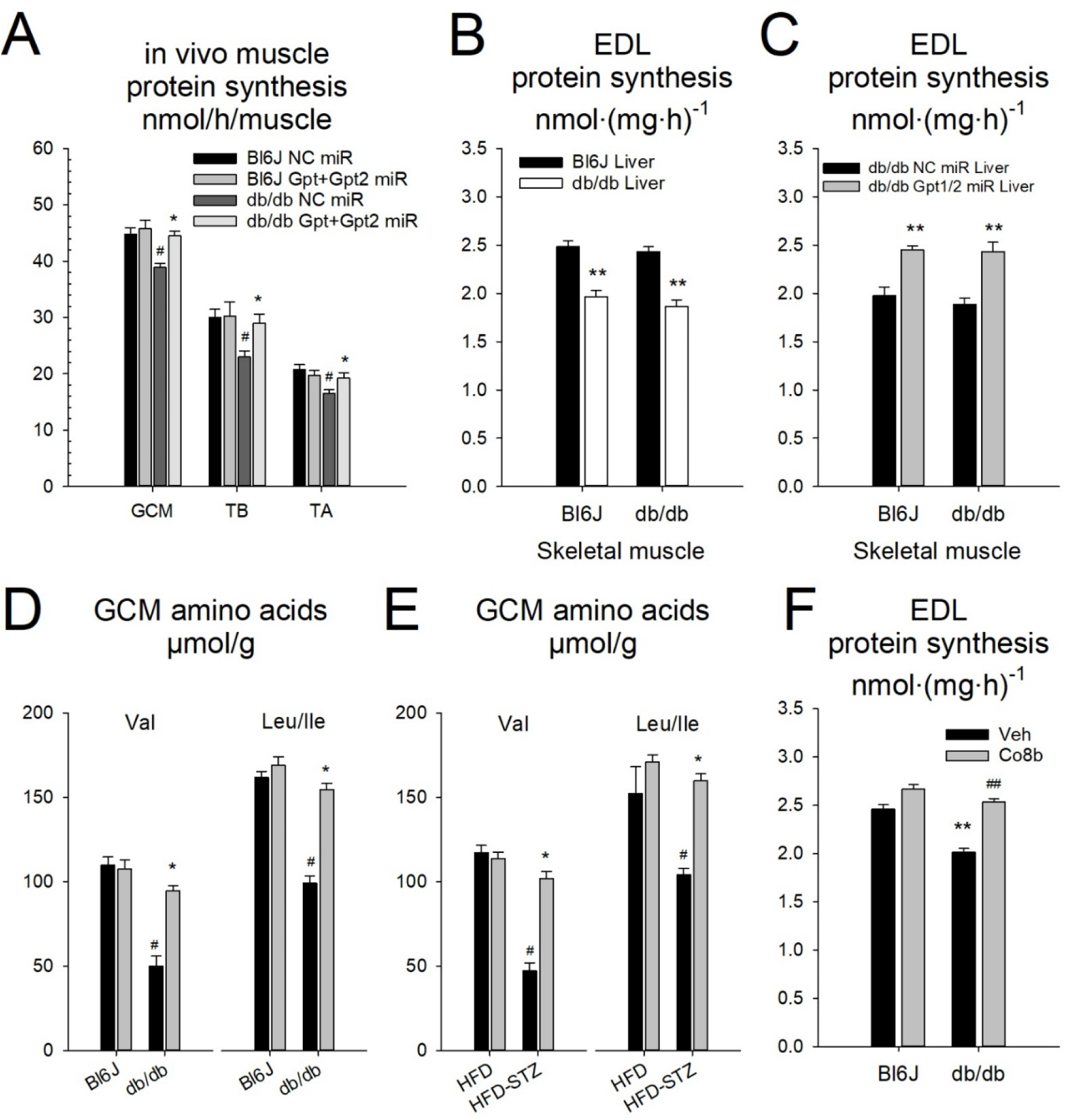
Disrupting the heightened liver alanine catabolism reverses the reduced skeletal muscle protein synthesis and branched chain amino acid levels in type 2 diabetes. A: In vivo protein synthesis rate calculated from mixed muscle ^3^H-phenylalanine incorporation in overnight fasted, 4h refed lean C57Bl/6J (Bl6) and age-matched obese/diabetic BKS-db/db mice with hepatocyte selective AAV-miR mediated silencing of glutamic-pyruvic transaminase (Gpt) isoforms. Study was conducted one week after AAV administrations. NC: negative control. miR: micro-RNA. GCM: gastrocnemius complex muscle. TB: Triceps brachii. TA: tibialis anterior. Data are mean ± SEM, N = 4/group. Effect of miR vs. NC miR: * P < 0.05, ** P < 0.01, *** P < 0.001. Effect of genotype: # P < 0.05, ## P < 0.01, ### P < 0.001. B: Ex vivo extensor digitorum longus (EDL) skeletal muscle protein synthesis rate during co-culture and cross-co-culture with liver slices. Tissues were taken from lean C57Bl/6J (Bl6) and age-matched obese/diabetic BKS-db/db mice with hepatocyte selective AAV-miR mediated silencing of glutamic-pyruvic transaminase (Gpt) isoforms. NC: negative control. miR: micro-RNA. Data are mean ± SEM, N = 3/group with 2 technical replicates per treatment condition. Effect of Liver genotype: * P < 0.05, ** P < 0.01, *** P < 0.001. Effect of muscle genotype: # P < 0.05, ## P < 0.01, ### P < 0.001. C: Ex vivo extensor digitorum longus (EDL) skeletal muscle protein synthesis rate during co-culture and cross-co-culture with liver slices. Tissues were taken from lean C57Bl/6J (Bl6) and age-matched obese/diabetic BKS-db/db mice with hepatocyte selective AAV-miR mediated silencing of glutamic-pyruvic transaminase (Gpt) isoforms. NC: negative control. miR: micro-RNA. Data are mean ± SEM, N = 3/group with 2 technical replicates per treatment condition. Effect of Liver miR: * P < 0.05, ** P < 0.01, *** P < 0.001. Effect of muscle genotype: # P < 0.05, ## P < 0.01, ### P < 0.001. D: Gastrocnemius complex skeletal muscle (GCM) valine (Val) and Leucine/Isoleucine (Leu/Ile) concentrations in lean C57Bl/6J (Bl6) and age-matched obese/diabetic BKS-db/db mice with hepatocyte selective AAV-miR mediated silencing of glutamic-pyruvic transaminase (Gpt) isoforms. NC: negative control. miR: micro-RNA. Data are mean ± SEM, N = 4/group. Effect of miR vs. NC miR: * P < 0.05, ** P < 0.01, *** P < 0.001. Effect of genotype: # P < 0.05, ## P < 0.01, ### P < 0.001. E: Gastrocnemius complex skeletal muscle (GCM) valine (Val) and Leucine/Isoleucine (Leu/Ile) concentrations in mice on an obesogenic high-fat diet with (HFD-STZ) or without (HFD) streptozotocin (STZ) pre-treatment to exacerbate the progression of frank diabetes; with hepatocyte selective AAV-miR mediated silencing of glutamic-pyruvic transaminase (Gpt) isoforms. NC: negative control. Data are mean ± SEM, N = 6/group. Effect of miR vs. NC miR: * P < 0.05, ** P < 0.01, *** P < 0.001. Effect of STZ: # P < 0.05, ## P < 0.01, ### P < 0.001. F: Ex vivo extensor digitorum longus (EDL) skeletal muscle protein synthesis rate during co-culture and cross-co-culture with liver slices with (compound 8b: Co8b) or without (Veh) treatment with an inhibitor of mitochondrial branched-chain amino acid transaminase. Tissues were taken from lean C57Bl/6J (Bl6) and age-matched obese/diabetic BKS-db/db mice. NC: negative control. miR: micro-RNA. Data are mean ± SEM, N = 3/group with 2 technical replicates per treatment condition. Effect of muscle genotype: * P < 0.05, ** P < 0.01, *** P < 0.001. Effect of Co8b: # P < 0.05, ## P < 0.01, ### P < 0.001. Statistical tests: A, B, C, E, F, G: 2-way ANOVA with Holm-Sidak posthoc hoc tests. D: 1-way ANOVA with Holm-Sidak posthoc hoc tests.

## Discussion

The major finding of the present studies was that we identified a liver-specific endocrine-metabolic mechanism linking hyperglycemia and muscle atrophy in diabetes. In particular, similar to prior studies ^41,42^ we demonstrated that liver alanine aminotransferase (ALT) isoform expression/activity is elevated in conditions of metabolic disease in mice and humans, and that liver-restricted ALT silencing *in vivo* retards, and reverses the progression of, hyperglycemia in mouse models of obesity and pre-as well as frank-type 2 diabetes (T2D). An unexpected result, however, was that the muscle atrophy of these mice was reversed upon liver ALT silencing. Importantly, obesity-driven metabolic disease and muscle atrophy are often linked in human T2D, the mechanisms behind which were poorly understood, and other than physical activity, no other therapeutic intervention thus far is efficacious in treating these two co-morbidities, at least to our knowledge. This finding is important as, other than metabolic control which is essential for prevention of other complications of diabetes, physical functional capability is linked to improved quality of life and late-life health ^2^.

The muscle atrophy, and related weakness, in T2D could be reversed by a silencing of the upregulation of liver alanine catabolism, an intriguing result. Such a result is not unfounded however, as some prior studies have linked liver metabolic control with systemic protein turnover and skeletal muscle phenotype ^43,44^. Similar to our studies in mice (Fig. 4), human individuals with T2D have lower muscle mass and strength ^45,46^, the mechanisms behind which are still not understood. It is known that hyperglycemia is associated with lower lean body mass in older humans ^47^, suggesting that there may be a metabolic basis of muscle atrophy. Indeed, T2D humans do have higher systemic protein and amino acid turnover across the whole body as well as in skeletal muscle, but do not exhibit lower muscle protein synthesis per se ^45,48,49^. Whereas some studies of obese/sarcopenic rats have demonstrated similar findings to the human studies ^50^, we (Fig. 7) and others ^37^ have demonstrated the some models of obese/diabetic mice such as the BKS-db/db mouse exhibits lower skeletal muscle protein synthesis, which is related to lower skeletal muscle size, fiber size, and strength. Our studies were conducted in the acute refed state and we express protein synthesis as a function of the entire muscle tissue, which may explain some of the differences with prior experiments. Interestingly, some, but not all, T2D preclinical mouse models exhibit hyperglucocorticoidemia, which we show to be an important factor linking obesity/T2D with sarcopenia (Fig. 6). Indeed, the Tallyho ^37^ and NZO (Fig. S5) obese/T2D mouse models have normal or higher skeletal muscle mass with correspondingly normal levels of corticosterone. This suggests that perhaps only a certain population of obese/T2D individuals develop muscle atrophy, and that this may be related to a hypercortisolism as humans with Cushing’s syndrome also exhibit sarcopenic obesity and T2D ^51^. Nevertheless, muscle branched-chain amino acid breakdown/turnover is greater in T2D humans ^52^, and we observed lower BCAA levels in obese/diabetic mouse skeletal muscle, suggesting that this may be a related factor. In particular, based upon cross-co-culture experiments with liver and muscle ex vivo (Fig. 5), we demonstrated that enhanced liver ALT activity in obesity/T2D directly controls skeletal muscle protein synthesis, perhaps via a pull mechanism to drain alanine from muscle and thereby enhance BCAA breakdown. BCAA catabolism can fuel both alanine release from skeletal muscle ^38^, and BCAA are key metabolites in the activation of mTORC1 ^53^, a key molecular control point for skeletal muscle size and protein synthesis ^54^. Even though it could be expected that a change in liver metabolism might act through a liver-secreted factor such as insulin-like growth factor 1, such hepatokines were not altered by liver ALT silencing in T2D mice (Fig. S7), experiments using dialyzed media from BKD-db/db liver (Fig. S7) discounted this possibility. However, ex vivo liver-muscle crosstalk experiments to blunt excess alanine expulsion from muscle such as media alanine excess or inhibiting muscle branched-chain amino acid transaminase (Fig. 7 and S7) supported a direct role for liver-muscle metabolic crosstalk in linking muscle atrophy and hyperglycemia in T2D.

Here we demonstrate that liver alanine catabolism by ALT functionally contributes to glycemic regulation under diverse nutritional conditions and the upregulation of ALT in obesity mice contributes to hyperglycemia and frank T2D. Importantly, enhanced splanchnic alanine clearance is also observed in obese and T2D humans ^13,14^ which is related to postprandial hyperglycemia upon protein feeding ^55^, suggesting that our pre-clinical studies are of clinical relevance for proper glucoregulation in humans with impaired metabolic control. The alanine aminotransferase isoforms, encoded by the genes *Gpt* and *Gpt2*, display a broad tissue expression profile and catalyze the reversible transamination of L-alanine and α-ketoglutarate to L-glutamate and pyruvate, with Gpt having a cytosolic- and Gpt2 a mitochondrial-localization ^56^. Despite its conception nearly 50 years ago ^11^, experimental evidence for the glucose-alanine cycle, which is a metabolic cycle involving the production of alanine from skeletal muscle and the production of glucose from alanine by the liver ^9^, remained obscure, despite some indirect indications ^57,58^. Concerning the contribution of ALT isoforms to systemic function, germline mutations in Gpt2 cause neurological deficiencies ^17,59^ presumably via the contribution of mitochondrial TCA cycle anaplerosis ^59^. Furthermore, similar to our results demonstrating the requirement of silencing both liver ALT isoforms for effects on systemic metabolism (Fig. 2 −4), *Gpt2* has been shown to be able to compensate for the loss of the mitochondrial pyruvate transporter in gluconeogenesis by the liver ^60,61^. Given that the contribution of liver ALT loss-of-function could only be revealed under specific conditions such as the direct provision of alanine or low-carbohydrate diets to force gluconeogenesis from amino acids, normoglycemia was remarkably well maintained because of adaptations in ketogenesis and probably gluconeogenesis from adipose triglyceride glycerol via the glycerol kinase reaction. Indeed, ex vivo hepatic glucose production was blunted in response to portal vein amino acids as well as alanine, but unresponsive to exogenous glycerol, with ALT isoform silencing (Fig. 2). This highlights the remarkable metabolic adaptability of the liver to utilize diverse substrates for specific metabolic outcomes, and adds to the existing knowledge of the relationship between amino acid and lipid metabolism ^12,62,63^.

Concerning T2D, liver ALT expression/activity is higher in T2D mice and humans (Fig. 1) and liver ALT silencing reduced hyperglycemia in the ad libitum fed state in mice (Fig. 3). Furthermore, ex vivo hepatic glucose production was exaggerated in response to portal vein amino acids as well as alanine in T2D, effects that were blunted with liver ALT silencing (Fig. 3), indicating that the effects of liver ALT silencing in vivo on glycaemia were cell-autonomous. It should be noted however, that liver ALT silencing did not completely blunt the hyperglycemia in our pre-clinical T2D studies, which is to be expected as there are several alternate sources of endogenous glucose production such as the kidney and intestine as well as several other sources of hepatic glucose production such as glycogenolysis and gluconeogenesis from lactate, glycerol as well as other glucogenic amino acids ^64^.

Lastly, we identified an hyperglucocorticoidemia-liver glucocorticoid receptor (GR) axis in T2D promoting heightened liver ALT isoform expression and systemic metabolic consequences thereof (Fig. 5). Glucocorticoids are known to have direct effects on skeletal muscle to induce muscle atrophy, despite intact liver GR signaling ^24,25^. Therefore, although the liver GR was not a critical component ^43^, our results demonstrate that blocking the previously observed ^65^ increase in liver ALT activity could confer an abrogation in the muscle atrophy upon synthetic glucocorticoid treatment. The lack of effects of the liver GR action might be explained by the partial action of liver-GR silencing on the upregulation of liver ALT expression (Fig. 6). Indeed, it is known that the majority of stress-pathway-induced genes in the liver require multiple inputs from different transcriptional nodes such as transcription factors downstream of glucagon signaling in addition to GR signaling ^66^. Moreover, glucagon signaling is known to affect expression of transcripts encoding amino acid catabolic enzymes ^67^, and glucocorticoid treatment may stimulate pancreatic islet glucagon production via increased circulating alanine levels ^26^. Importantly, glucagon levels are higher in humans ^31^ and mice (Fig. 6) with T2D, and co-treatment with glucocorticoid and glucagon further enhanced liver Gpt/2 expression (Fig. S6). Despite glucagon receptor not being expressed in skeletal muscle but with potent liver expression and signaling (Fig. S6), chronic glucagon levels promote lean tissue wasting ^29^ and glucagonoma patients are known to present with muscle weakness ^68^. Given that liver Gpt/2 silencing prevented skeletal muscle wasting and hyperglycaemia with chronic glucagon treatment (Fig. 6), taken altogether this then exemplifies the role of an endocrine-liver-muscle axis in linking muscle atrophy and hyperglycaemia in T2D.

Taken together, the results of our preclinical studies provide intriguing new evidence for an endocrine-liver-amino acid catabolic axis that links both hyperglycemia and muscle atrophy in T2D.

## Methods

### Mouse experiments

Unless stated otherwise, male mice aged 7 weeks upon arrival, were acclimated to the local housing facility (12-12h light-dark cycle, 22-24°C) for one week prior to experimentation and were fed standard rodent chow (3437, Provimi Kliba, DEU) unless otherwise indicated. Mice used for experiments were C57Bl/6J (#000664, Charles River Laboratories or Monash Anmal Research Platform, AUS) C57Bl/6NCrl mice (#027, Charles River Laboratories), as well as BKS.Cg-Dock7^m^+/+Lepr^db^J (BKS-db/db; #000642, Charles River Laboratories). Samples from prior mouse experiments analysed here include: 9-week old male pre-diabetic obese ob/ob mice ^69^; 8-week old male New Zealand Black (NZB) and obese/pre-diabetic New Zealand Obese (NZO) as well as 12 week old C57Bl/6J and BKS-db/db subjected 24-h fasting and refeeding regimes ^15^; 16 week low-fat or high-fat diet feeding in C57Bl/6N mice from 8 weeks of age as well as NZB and diabetic NZO mice kept on a low-fat diet from 8 until 18 weeks of age ^19^; glucocorticoid receptor (GR; Nr3c1) floxed/floxed C57Bl/6J mice administered negative control or CRE-expressing adenoviruses an then subjected to dexamethasone treatment ^43^; BKS-db/db mice were administered negative control or GR-specific shRNA expression adenoviruses and sacrificed one week later ^70^.

For liver-specific Gpt and Gpt2 silencing or overexpression experiments, we conducted experiments where following acclimation, mice were administered a total of 2 × 10^11^ virus particles per mouse via the tail vein. For the negative control (NC): 2 × 10^11^ NC miR-AAV; Gpt: 1 × 10^11^ Gpt miR-AAV + 1 × 10^11^ NC miR-AAV; Gpt2: 1 × 10^11^ Gpt2 miR-AAV + 1 × 10^11^ NC miR-AAV; Gpt+Gpt2: 1 × 10^11^ Gpt miR-AAV + 1 × 10^11^ Gpt2 miR-AAV. For overexpression experiments, in addition to the NC and Gpt+Gpt2, mice were administered 1 × 10^11^ hsGpt(wt) cDNA + 1 × 10^11^ hsGpt2(wt) cDNA or 1 × 10^11^ hsGpt(mut) cDNA + 1 × 10^11^ hsGpt2(mut) cDNA. Human Flag- and myc-tagged GPT and GPT2 cDNA was obtained from Origene (RC203756 and RC209119). Mutagenesis of Ser126 to Arg in GPT and Ser153 to ARg in GPT2 was performed with the Q5 site-directed mutagenesis kit frm NEB (#E0554).

To initially assess the effects of liver Gpt/Gpt2 silencing, we administered the AAVs to C57Bl/6N mice, after which we allowed the transduction and silencing to take place for 2 weeks. Then we conducted a battery of metabolic phenotyping experiments over a further 4 week period. Furthermore, in follow-up experiments mice were subjected to either ketogenic (i.e. 89.5%E fat, 10.4%E protein; D12369B, Research Diets, USA) or protein-enriched (80%E protein, 10% carbohydrate, 10% fat; D16080303, Research Diets, USA) diets for a period of 3 weeks with various metabolic phenotyping tests. To assess the effects of liver Gpt/Gpt2 silencing in mouse models of diabetes, we initially conducted a study in 12 week old BKS-db/db mice where we administered AAV and two weeks later conducted a battery of phenotyping experiments over a further 6 week period. We the followed up this study by conducted a strain comparison study, which focussed on skeletal muscle, where we examined the effects of liver Gpt/Gpt2 silencing on C57Bl/6J and BKS-db/db mice where viruses were administered at 8 weeks of age and with phenotyping experiment then conducted between 10-16 weeks of age after which time the experiment was terminated. To assess the effects of liver Gpt/Gpt2 in an alternate mouse model of diabetes, we employed the high-fat diet with low-dose streptozotocin model essentially as described ^71,72^. To conduct this, we administered AAVs to 7 week old C57Bl/6N mice, 3d after which we placed all mice on a high-fat diet (i.e. 60%E fat, D12492, Research Diets) for 7d, after which half of the mice in each AAV group received 25mg/kg streptozotocin (S7870, LKT laboratories Inc., USA) or vehicle (Na-citrate buffer, pH 4.5) for 3 consecutive days. The mice were then maintained on the HFD for a further 11 weeks after which time the experiment was terminated. To test the involvement of liver Gpt/Gpt2 in systemic glucocorticoid exposure, we administered AAVs to 8 week old C57Bl/6N mice 2 weeks after which we injected intraperitoneally on a daily basis with 9α-Fluoro-16α-methylprednisolone (Dexamethasone, DEX; 1 mg/kg, D8893 Sigma–Aldrich, DEU) or Vehicle (VEH, 2% ethanol in isotonic saline) for 21d. By way of follow up, we then employed high-fat diet feeding together with chronic glucocorticoid treatment, as strategy known to produce severe metabolic dysfunction in rodents ^20–22^. For this, we administered AAVs to 8 week old C57Bl/6N mice 2 weeks after which we initiated feeding of a high fat diet (60% calories from fat; SF13-092, Specialty Feeds, AUS) and injected DEX intraperitoneally (1mg/kg; twice weekly) for 6 weeks. Lastly, similar to a prior study ^32^, we conducted a study where a stable glucagon peptide (i.e. acyl-GCG; synthesized by Auspep (AUS) according ^32^) to was administered to mice. Specifically, we administered AAVs to 8 week old C57Bl/6N mice, 2 weeks after which we intraperitoneally administered acyl-GCG (10nmol/kg/d; IUB288 (batch BF20342), Auspep, AUS) for 21d. For acute acyl-GCG experiments, a single dose of acyl-GCG (10nmol/kg/d) was administered by IP injection after a 4h fast.

For glucagon receptor (GCGR) inhibition experiments, a human monoclonal GCGR antibody (REMD), known to have potent effects in T2D mice to reduce blood glucose ^73^, was administed to C57Bl/6J and BKS-db/db mice (Jackson Laboratories, USA). More specifically, 6 wk old mice were administered were administered REMD (5mg/kg/d) or vehicle (PBS) for 7d after which they were sacrificed and liver tissue was harvested and snap frozen in LN2 prior to stargaze at −80°C.

For all studies, phenotyping tests were separated by at least one week and mice were sacrificed between ZT3-6 in the ad libitum fed state and tissues including blood serum, urine, liver, perigonadal fat and gastrocnemius muscles were carefully and rapidly excised, weighed precisely, and then frozen in LN2 or placed in fixation solution (P087.5, Roth GmbH, DEU) for histological assessment. In some experiments, tibialis anterior and biceps brachii skeletal muscles were also excised and weighed. Animal experiments were conducted according to regional, national, and continental ethical guidelines and protocols were approved by local regulatory authorities (Regierungspräsidium Karlsruhe, DEU; Monash University Animal Ethics Committee, AUS) and conformed to ARRIVE guidelines.

### Human subjects

Gene transcript expression was investigated in liver tissue samples obtained from 10 Caucasian healthy donors and 64 Caucasian obese men and women with (*n* = 27) or without (*n* = 37) type 2 diabetes who underwent open abdominal surgery for Roux-en-Y bypass, sleeve gastrectomy or elective cholecystectomy. A small liver biopsy was taken during the surgery, immediately frozen in liquid nitrogen, and stored at −80°C until further preparations. The phenotypic characterization of the cohort has been performed as described previously ^74^. Serum samples and liver biopsies were taken between 8 am and 10 am after an overnight fast. For the human experiments, the study was approved by the local ethics committee of the University of Leipzig, Germany (363-10-13122010 and 017-12-230112). All patients gave preoperative written informed consent for the use of their samples.

### Recombinant viruses

For miRNA experiments, oligonucleotides targeting mouse *Gpt* (5′-AATATATTGCGCCACATCCTC-3′) and *Gpt2* (5′-TTCACAACAATGTCCATGGCG-3′) as well as non-specific negative control oligonucleotides (5′-AAATGTACTGCGCGTGGAGAC-3′) were cloned into pcDNA6.2-GW/EmGFP-miR [“BLOCK-iT™ PolII miR RNAi Expression Vector Kit” (Invitrogen, Darmstadt, DEU). Human Flag- and myc-tagged GPT and GPT2 cDNA was obtained from Origene (RC203756 and RC209119). Mutagenesis of Ser125 to Arg in GPT and Ser153 to Arg in GPT2 ^17^ was performed with the Q5 site-directed mutagenesis kit from NEB (#E0554). Adeno-associated viruses (AAV) encoding control or specific miRNAs or GPT/2 cDNAs under the control of a hepatocyte-specific promoter were established, purified and titered as described previously ^16,75^. In particular, AAV vectors were purified using iodixanol step gradients and titrated as previously described in detail ^76^.

### Biometric, metabolic and functional phenotyping

#### Biometric and functional measurements

In all experiments mice were carefully weighed on a precise scale every week. In some experiments, 2d prior to sacrifice, body composition was assessed by ECHO-magnetic resonance imaging (Echo Medical Systems, USA) with an initial measurement taken in some cases. In some experiments, neuromuscular function was assessed immediately before ECHO-MRI analysis using a grip strength tester (BIO-GS3, BIOSEB, France) with the average of three measurements taken per mouse.

#### Dynamic metabolic tests in vivo

In some experiments we conducted a fasting-feeding regimen. This involved placing the mice in a new cage without food but with water overnight, followed by 6h of refeeding. In some experiments we conducted an alanine tolerance test, whereby mice were fasted overnight, and subsequently administered by intraperitoneal injection, a fixed dose of 25mg alanine (i.e. 250µL of a 10% solution in 0.9% saline). Blood samples were taken from the tail vein before and at defined time points after the alanine administration. In all experiments, blood samples taken from the tail vein before and at defined time points after the challenge for blood glucose measurement (AccuCheck Aviva) as well as collection in heparinized tubes (Microvette CB 300 LH, Sarstedt) for eventual blood plasma analyses.

#### In vivo protein synthesis

To conduct this we adapted a method described ^77^. In the ad libitum fed state between ZT3-ZT7, mice were injected with L-[2,3,4,5,6-^3^H]phenylalanine (NET112200, Perkin Elmer; 150 mM, 30 µCi/ml; 0.5 mL], with blood samples taken from the tail vein at time 0, 4, 8 and 12 min with the mice sacrificed at 15 min and skeletal muscle tissues were rapidly harvested. Muscles were homogenized and protein precipitated and re-dissolved, with protein extracts and serum samples then subjected to liquid-scintillation counting (Packard 2200CA Tri-Carb Liquid Scintillation analyzer; Packard Instruments). Tissue uptake rates were calculated based upon the weighted average of the specific activity of the administered ^3^H-Phe (cpm/mol), along with tissue counts (cpm) and mass (g), and time (h) of collection after administration.

#### Ex vivo skeletal muscle protein synthesis with liver slice co-culture

Extensor digitorum longus (EDL) muscles and precision cut sliver slices were dissected from C57Bl/6J or BKS-db/db mice and co-cultured in non-adherent 24 well tissue culture plates (662102, Greiner Bio-One) with inserts for co-culture (665641, Greiner Bio-One). In brief, these experiments were carried out on precision-cut liver slices as previously described ^78^. In particular, liver slices and EDL muscles were taken and prepared in UW solution (in absence of Insulin and Dexamethasone) and then pre-incubated for 30min in Williams media E containing 50 μg/mL gentamycin, 5% dialyzed FBS, 11 mM d-glucose, 0.3 mM pyruvate, 0.1 μM methyl-linoleate, and an amino acid mixture resembling the hepatic portal vein (PVAA) concentrations ^79^ formulated from custom made 2× William’s Media E without macro-nutrients (C4318.0500SPI, Genaxxon Bioscience GmbH). After this, media was replaced with media plus tracer (i.e. L-^14^C-U-Leucine, NEC279E0, Perkin Elmer) plus 10 nM insulin (91077C, Sigma-Aldrich) and 10 nM glucagon (G2044, Sigma-Aldrich) and then incubated for 3h after which muscles were washed twice in ice cold PBS, carefully weighed and frozen. To determine protein synthesis ^80^, muscles were then lysed in 1.5mL of lysis buffer and tricholoacetic acid precipitated proteins were then re-solubilized and radioactive leucine incorporation into the total protein pool was assessed by liquid scintillation counting. Protein synthesis rate was calculated from specific activity of the media leucine (cpm/mol), along with tissue protein counts (cpm) and mass (g), and time (h) of the tracer assay.

Experiments were conducted with different combinations of C57Bl/6J and BKS-db/db liver slices and skeletal muscles with or without prior AAV-mediated Gpt/Gpt2 silencing. Additional experiments were conducted where media was first incubated with BKS-db/db liver for 3h and then dialyzed (1 kDa cutoff, GE80-6483-75, Sigma, DEU) at 4°C against fresh Williams media E (formulation indicated above) to revert any metabolite changes but leave larger molecules such as peptides present. This dialyzed media was then immediately used to assess effects on protein synthesis in isolated C57Bl/6J EDL muscles. In addition, studies were conducted where 5mM alanine (Sigma, DEU) or 20µM compound 8b (Co8b; a BCAT2 inhibitor, pIC50 Cell 6.2 ^40^; MU-002-BC19 (batch KPS-266-45e) Advanced Molecular Technologies, AUS) in the co-incubation media.

#### Ex vivo liver glucose production

We conducted these studies essentially according to ^15^ with minor adaptations. In brief, liver slices were prepared as above from overnight fasted C57Bl/6J or BKS-db/db mice and then incubated with a control media (Williams media E containing 50 μg/mL gentamycin, 5% dialyzed FBS, 0mM d-glucose, 0.03mM pyruvate, 0.1 μM methyl-linoleate, 0.05X PVAA), as well as control media with 1X PVAA, 1mM alanine, 1mM glutamine, 1 mM pyruvate, or 1mM glycerol for 18h. Glucose concentration of the media was measured (GAHK20, Sigma-Aldrich, DEU) and subsequently glucose production rate was calculated from media concentrations from liver slice incubations minus those from media without slice incubations, divided by the time of incubation (i.e. (slice media (M) – control media (M))/time (h)). Importantly, liver slice weights (4.4 ± 0.3 mg) did not differ between any of the conditions studied.

#### Blood and urine metabolites & hormones

Blood glucose was measured using an automated analyzer (AccuCheck Aviva). Commercially available kits were used to measure serum non-esterified fatty acids (NEFA; NEFA-HR, Wako, USA), glycerol/triglyceride (TG; TR-0100; Sigma-Aldrich, DEU), cholesterol (CH200, Randox, USA), ketone bodies (415-73301, T-KB, Wako, USA), lactate (K607-100; Biovision), and corticosterone (900-097, Assay Designs, USA) essentially according to manufacturer’s instructions. Urine glucose (GAHK20, Sigma-Aldrich, DEU) and creatinine (K625-100, Biovision, USA) analyses were conducted using commercially available kits. All samples were loaded in order to fit within the assay range of the reagents supplied.

Amino acids were determined in serum/plasma by electrospray ionization tandem mass spectrometry (ESI-MS/MS) according to a modified method as previously described ^81^, using a Quattro Ultima triple quadrupole mass spectrometer (Micromass, Manchester, UK) equipped with an electrospray ion source and a Micromass MassLynx data system. For tissue determinations, frozen tissue was carefully weighed and then extracted in 10 volumes of ice cold PBS. Values obtained were then calculated to give μmol/g wet tissue weight.

#### RNA extraction and analysis

Total RNA was extracted from homogenized mouse tissue using TRIzol reagent (Life Technologies, DEU). cDNA was prepared by reverse transcription using the Quantitect RT-PCR kit (205313, Qiagen, DEU). cDNAs were amplified using assay-on-demand kits (*Gpt*: Mm00805379_g1; *GPT*: Hs00193397_m1; *Gpt2*: Mm00558028_m1; *GPT2*: Hs00370287_m1; *Tbp*: Mm01277042_m1; *HPRT1*: Hs01003267_m1) and an ABIPRISM 7700 Sequence detector (Applied Biosystems, Darmstadt, DEU). RNA expression data was normalized to levels of *TATA-box binding protein* (*Tbp;* mouse) or *hypoxanthine guanine phosphoribosyltransferase 1* (*HPRT1*; human) mRNA.

#### Tissue protein expression analysis

For simple expression analyses from homogenates, protein was extracted (Qiagen Tissue Lyser) from ∼50 mg frozen liver in tissue lysis buffer ^80^. To determine protein expression, total protein concentration was analyzed by BCA assay (23227, Fischer Scientific, DEU) and equal amounts of denatured protein were loaded onto an SDS-polyacrylamide gel, subjected to electrophoresis and blotted onto nitrocellulose membrane. Western blot assays were performed using standard techniques with antibodies specific for with antibodies against GPT (sc-47017, Santa-Cruz Biotech; 1:1000), GPT2 (AP18028a, Abgent; 1:1000), and the housekeeping protein HSP90 (610418, BD Biosciences; 1:5000) and chemiluminescent images were captured using a ChemiDoc imaging system with ImageLab software (Bio-Rad). For expression analyses, all western blots signals were within a linear range, with respect to protein loaded and antibodies used, as determined by preliminary experiments, and detected bands ran at the expected relative mobility (data not shown).

#### Tissue ALT assay

Tissue ALT activity was determined using a commercially available kit following the kit instructions (K752, Biovision, USA). Briefly, ~50mg of liver tissue was homogenized (Qiagen Tissue Lyser) while cold in ALT lysis buffer and 10uL of extract with 3 technical replicates used. Assay data were normalized to total extract protein in 10uL as determined by a total protein assay (23227, Fischer Scientific, DEU).

#### Tissue histology

For *GPT* and *GPT2* staining, dduring liver dissection a piece of liver was carefully dissected and then fixed in 4% formalin (P087.5, Roth GmbH, DEU), washed and subsequently dehydrated and embedded in paraffin. 2µM paraffin sections were cut, followed by antigen retrieval with EDTA buffer, after which they were incubated with antibodies against GPT (AP7468b, Abgent; 1:200) and GPT2 (PAD854Mu01, Cloud-Clone Inc.; 1:200) antigens in BOND^TM^ primary antibody diluent (AR9352, Leica Biosystems). For skeletal muscle fiber size visualization, whole gastrocnemius muscles were carefully dissected and then fixed in 4% formalin (P087.5, Roth GmbH, DEU), washed and subsequently dehydrated and embedded in paraffin. 2µM paraffin cross-sections were cut, followed by antigen retrieval with citrate buffer, after which they were incubated with antibodies against dystrophin (ab15277, Abcam; 1:500) in BOND^TM^ primary antibody diluent (AR9352, Leica Biosystems). In both cases, primary antibody exposure was followed by secondary antibody (Leica Biosystems) and staining using the Bond^TM^ Polymer Refine Detection kit (DS9800, Leica Biosystems). Brightfield images were acquired using a Leica DM 1000 LED microscope and processed using Leica Application Suite software (Leica Biosystems).

#### Gpt and Gpt2 gene transcriptional regulation

Publically available sequencing data ^23^ were analyzed in silico in order to ascertain DEX-dependent GR-DNA binding regions. Accessible NCBI Gene Expression Omnibus GSE files for GR & CEBPB Chip-Seqs and DNAse hypersensitivity were downloaded and uploaded to Ensembl Mouse Genome Browser NCBI m37. The browser was then configured to display *Gpt* and *Gpt2* gene and promoter regions, as well as flanking regions, and images were taken.

### Quantification and statistical analyses

#### Statistical analyses

Mice were assigned to groups based upon initial body mass for counterbalancing. Pre-established criteria for exclusion of mice from study groups were obvious infections/wounds which would impact on feeding behavior as well as metabolic profile. Where possible, analysis of samples was blinded.

Statistical analyses were performed using t-tests (two-sided), or 2-way analysis of variance (ANOVA) with or without repeated measures, where appropriate, with Holm-Sidak-adjusted post-tests. Nonparametric tests (e.g. Mann–Whitney–Wilcoxon) were conducted when data were not normally distributed. All analyses were carried out with SigmaPlot v.13 software (Systat Software GmbH, DEU). Statistical details can be found within the Figure legends. Differences between groups were considered significant when P < 0.05. Raw data are available upon request.

The authors have declared that no conflict of interest exists.

## Author contributions

Project conceptualization, administration and management: AJR. Resources: JGO, MB, SH, OM, MH, AJR. Investigation: JGO, JS, KVS, AZ, AJ, LM, MB, MH, AJR. Software and formal analysis: PR, AJR. Writing, original draft: AJR. Writing, editing: JGO, PMR, JS, MB, SH, OM, MH, AJR. Visualization: MH, AJR. Funding acquisition: SH, MH, AJR.

## Acknowledgements

The authors wish to thank Jessica Fuhrmeister, Adriano Maida and Thomas Gantert (DKFZ, A170), Lena Figur (A171, DKFZ), as well as Jenny Hetzer and Danijela Heide (F180, DKFZ) for experimental support. JS was supported by a fellowship from the Helmholtz International Graduate School for Cancer Research. MH was supported by an ERC Consolidator grant (HepatoMetaboPath), SFBTR 209 (Liver Cancer), and SFBTR179. This project has received funding from the European Union’s Horizon 2020 research and innovation programme under grant agreement No 667273. This work was funded by the Helmholtz Future Topic “Aging and metabolic reprogramming” to SH and and by a project grant from the EFSD/Lilly European Diabetes Research Programme to AJR.

