## Supplementary material for "An endocrine-hepato-muscular metabolic cycle links skeletal muscle atrophy and hyperglycemia in type 2 diabetes": Supp file

**Supplemental Information**

Contents

Table S1

Table S2

Table S3

Figure S1

Figure S2

Figure S3

Figure S4

Figure S5

Figure S6

Figure S7

**Table S1.** Blood serum amino acid levels from lean and obese/diabetic mice in different nutritional states. NZB: New Zealand Black. NZO: New Zealand Obese. Data are mean ± SEM. Related to Fig. S1.

| Amino acid | C57Bl/6J | | | BKS-db/db | | |
|---|---|---|---|---|---|---|
| | *ad libitum fed* | *fasted* | *refed* | *ad libitum fed* | *fasted* | *refed* |
| **Ala** | 383 ± 41 | 406 ± 126 | 838 ± 77 | 445 ± 83 | 150 ± 9 | 467 ± 21 |
| **Pro** | 190 ± 15 | 263 ± 86 | 311 ± 82 | 243 ± 12 | 206 ± 15 | 198 ± 17 |
| **Val** | 286 ± 31 | 613 ± 235 | 682 ± 211 | 286 ± 19 | 279 ± 19 | 250 ± 15 |
| **Thr** | 108 ± 21 | 110 ± 42 | 157 ± 35 | 51 ± 3 | 31 ± 3 | 86 ± 7 |
| **Leu/Ile** | 94 ± 5 | 205 ± 71 | 221 ± 54 | 104 ± 5 | 96 ± 3 | 97 ± 1 |
| **Gln** | 113 ± 15 | 212 ± 81 | 242 ± 57 | 85 ± 5 | 74 ± 4 | 110 ± 15 |
| **Met** | 104 ± 16 | 246 ± 108 | 333 ± 81 | 68 ± 6 | 44 ± 3 | 108 ± 10 |
| **His** | 462 ± 42 | 567 ± 142 | 676 ± 117 | 439 ± 86 | 266 ± 19 | 374 ± 29 |
| **Phe** | 7,06 ± 0,67 | 6,91 ± 1,69 | 12,60 ± 1,15 | 9,09 ± 2,30 | 7,32 ± 0,92 | 8,34 ± 1,11 |
| **Tyr** | 94 ± 6 | 136 ± 44 | 182 ± 31 | 93 ± 4 | 97 ± 6 | 86 ± 9 |
| **Asx** | 282 ± 38 | 451 ± 196 | 657 ± 149 | 178 ± 22 | 72 ± 5 | 273 ± 20 |
| **Glx** | 682 ± 77 | 1339 ± 129 | 588 ± 99 | 135 ± 41 | 891 ± 91 | 136 ± 18 |
| **Trp** | 275 ± 27 | 373 ± 126 | 586 ± 100 | 180 ± 19 | 100 ± 6 | 266 ± 20 |
| **Gly** | 503 ± 30 | 501 ± 129 | 674 ± 38 | 311 ± 49 | 185 ± 10 | 314 ± 11 |
| **Orn** | 458 ± 52 | 363 ± 99 | 474 ± 27 | 863 ± 46 | 401 ± 34 | 503 ± 22 |
| **Arg** | 751 ± 96 | 1471 ± 582 | 1691 ± 468 | 155 ± 62 | 278 ± 48 | 762 ± 89 |
| **Cit** | 76 ± 4 | 53 ± 6 | 76 ± 2 | 104 ± 7 | 76 ± 5 | 107 ± 4 |
| **Hci** | 0,50 ± 0,07 | 1,07 ± 0,47 | 1,27 ± 0,34 | 0,43 ± 0,02 | 0,26 ± 0,01 | 0,55 ± 0,04 |
| **Asa** | 10,3 ± 0,7 | 7,5 ± 0,7 | 11,7 ± 0,6 | 9,8 ± 0,4 | 7,2 ± 0,6 | 9,8 ± 0,7 |

| Amino acid | NZB | | NZO | |
|---|---|---|---|---|
| | *fasted* | *refed* | *fasted* | *refed* |
| **Ala** | 709 ± 117 | 1398 ± 100 | 2501 ± 343 | 906 ± 116 |
| **Pro** | 543 ± 90 | 400 ± 47 | 1603 ± 239 | 522 ± 128 |
| **Val** | 1033 ± 250 | 649 ± 12 | 3871 ± 739 | 961 ± 285 |
| **Thr** | 354 ± 62 | 214 ± 17 | 493 ± 123 | 570 ± 228 |
| **Leu/Ile** | 274 ± 77 | 219 ± 5 | 1398 ± 399 | 355 ± 101 |
| **Gln** | 249 ± 90 | 265 ± 23 | 1259 ± 334 | 408 ± 108 |
| **Met** | 336 ± 85 | 214 ± 26 | 1481 ± 392 | 477 ± 162 |
| **His** | 1487 ± 287 | 1100 ± 108 | 3126 ± 649 | 1324 ± 376 |
| **Phe** | 81,6 ± 25,3 | 38,0 ± 3,2 | 39,5 ± 6,9 | 53,6 ± 9,7 |
| **Tyr** | 313 ± 60 | 279 ± 25 | 752 ± 169 | 289 ± 68 |
| **Asx** | 540 ± 180 | 725 ± 76 | 2414 ± 634 | 776 ± 287 |
| **Glx** | 809 ± 186 | 624 ± 43 | 2299 ± 653 | 3878 ± 2314 |
| **Trp** | 268 ± 63 | 252 ± 25 | 773 ± 174 | 297 ± 79 |
| **Gly** | 386 ± 62 | 319 ± 19 | 570 ± 82 | 266 ± 44 |
| **Orn** | 471 ± 74 | 509 ± 135 | 464 ± 52 | 366 ± 90 |
| **Arg** | 364 ± 206 | 289 ± 52 | 2332 ± 519 | 789 ± 297 |
| **Cit** | 52 ± 3 | 93 ± 2 | 60 ± 2 | 81 ± 7 |
| **Hci** | 1,70 ± 0,37 | 1,92 ± 0,32 | 7,68 ± 1,99 | 1,88 ± 0,70 |
| **Asa** | 5,9 ± 0,5 | 8,2 ± 0,3 | 8,3 ± 0,7 | 10,3 ± 0,9 |

**Table S2. Subject characteristics of obese type 2 diabetic (T2D) and normal glucose tolerant (NGT) obese subjects.** Parameters included body mass index (BMI), waist-to-hip-ratio (WHR), visceral (Vis), subcutaneous (SC) and total fat area, glycated haemoglobin (HbA1c), fasting plasma glucose (FP) and insulin (FPI), homeostatic model assessment of insulin resistance (HOMA-IR) and beta-cell function (HOMA-B), quantitative insulin sensitivity check index (QUICKI), clamp glucose infusion rate (GIR), total (total Chol) and LDL cholesterol (LDL-Chol), and triglycerides (TG). Data are mean ± SEM. Related to Fig. 1.

| | **NGT** | **T2D** | p-value |
|---|---|---|---|
| **M/F** | 23/17 | 14/11 | |
| **Height (m)** | 1.73 ± 0.01 | 1.72 ± 0.02 | 0.7095 |
| **Weight (kg)** | 93.6 ± 3.1 | 99.8 ± 4.63 | 0.2712 |
| **BMI (kg/m²)** | 31.3 ± 1.1 | 33.5 ± 1.2 | 0.1929 |
| **WHR** | 1.02 ± 0.02 | 1.05 ± 0.02 | 0.2631 |
| **Vis fat area (cm²)** | 151 ± 14 | 216 ± 18 | 0.0058 |
| **SC fat area (cm²)** | 549 ± 64 | 628 ± 82 | 0.4467 |
| **Total fat area (cm²)** | 700 ± 71 | 844 ± 82 | 0.1868 |
| **Liver fat (%)** | 14.2 ± 2.0 | 20.2 ± 3.0 | 0.0988 |
| **HbA1c (%)** | 5.52 ± 0.04 | 6.10 ± 0.11 | 0.0000 |
| **FPG (mM)** | 5.25 ± 0.07 | 6.42 ± 0.14 | 0.0000 |
| **FPI (pM)** | 93.3 ± 13.3 | 289.1 ± 25.3 | 0.0000 |
| **HOMA-IR** | 3.14 ± 0.46 | 11.98 ± 1.22 | 0.0000 |
| **HOMA-B** | 1.68 ± 0.25 | 2.98 ± 0.27 | 0.0009 |
| **QUICKI** | 0.362 ± 0.013 | 0.276 ± 0.003 | 0.0000 |
| **Clamp GIR (µmol/kg/min)** | 71.7 ± 4.5 | 36.9 ± 4.2 | 0.0000 |
| **Total Chol (mM)** | 4.86 ± 0.12 | 5.74 ± 0.16 | 0.0001 |
| **LDL-Chol (mM)** | 2.74 ± 0.12 | 3.56 ± 0.16 | 0.0002 |
| **TG (mM)** | 1.53 ± 0.11 | 2.38 ± 0.12 | 0.0000 |
| **Liver *GPT* mRNA (AU)** | 1.33 ± 0.20 | 4.08 ± 0.59 | 0.0001 |
| **Liver *GPT2* mRNA (AU)** | 0.56 ± 0.05 | 3.62 ± 0.67 | 0.0001 |

**Table S3. Correlative analysis of liver glutamic-pyruvic transaminase (*GPT*) and glutamic-pyruvic transaminase 2 (*GPT2*) mRNA expression and various biometric and serological parameters**. Parameters included body mass index (BMI), visceral (Vis), subcutaneous (SC) and total fat area, glycated haemoglobin (HbA1c), fasting plasma glucose (FP) and insulin (FPI), homeostatic model assessment of insulin resitance (HOMA-IR) and beta-cell function (HOMA-B), quantitative insulin sensitivity check index (QUICKI), clamp glucose infusion rate (GIR), total (total Chol) and LDL cholesterol (LDL-Chol), and triglycerides (TG). Shown are the Pearson's r product moment and associated p-value. Related to Fig. 1.

| | ***GPT*** | | ***GPT2*** | |
|---|---|---|---|---|
| | **Pearson's r** | **p** | **Pearson's r** | **p** |
| **Weight (kg)** | 0.181 | 0.1460 | 0.127 | 0.3090 |
| **BMI (kg/m2)** | 0.199 | 0.1100 | 0.131 | 0.2930 |
| **WHR** | 0.261 | 0.0327 | 0.303 | 0.0127 |
| **Vis fat area (cm²)** | 0.552 | 0.0000 | 0.268 | 0.0296 |
| **SC fat area (cm²)** | 0.0699 | 0.5770 | 0.151 | 0.2250 |
| **Total fat area (cm²)** | 0.183 | 0.1410 | 0.198 | 0.1110 |
| **Liver fat (%)** | 0.434 | 0.0003 | 0.194 | 0.1190 |
| **HbA1c (%)** | 0.268 | 0.0294 | 0.429 | 0.0003 |
| **FPG (mM)** | 0.275 | 0.0256 | 0.481 | 0.0000 |
| **FPI (pM)** | 0.45 | 0.0001 | 0.305 | 0.0128 |
| **HOMA-IR** | 0.42 | 0.0004 | 0.325 | 0.0077 |
| **HOMA-B** | 0.383 | 0.0015 | 0.114 | 0.3640 |
| **QUICKI** | -0.378 | 0.0018 | -0.311 | 0.0110 |
| **Clamp GIR (µmol/kg/min)** | -0.405 | 0.0007 | -0.27 | 0.0281 |
| **Total Chol (mM)** | -0.0123 | 0.9220 | 0.351 | 0.0039 |
| **LDL-Chol (mM)** | 0.382 | 0.0015 | 0.369 | 0.0023 |
| **TG (mM)** | 0.331 | 0.0063 | 0.296 | 0.0159 |
| **Cortisol (nM)** | 0.336 | 0.0093 | 0.258 | 0.0449 |

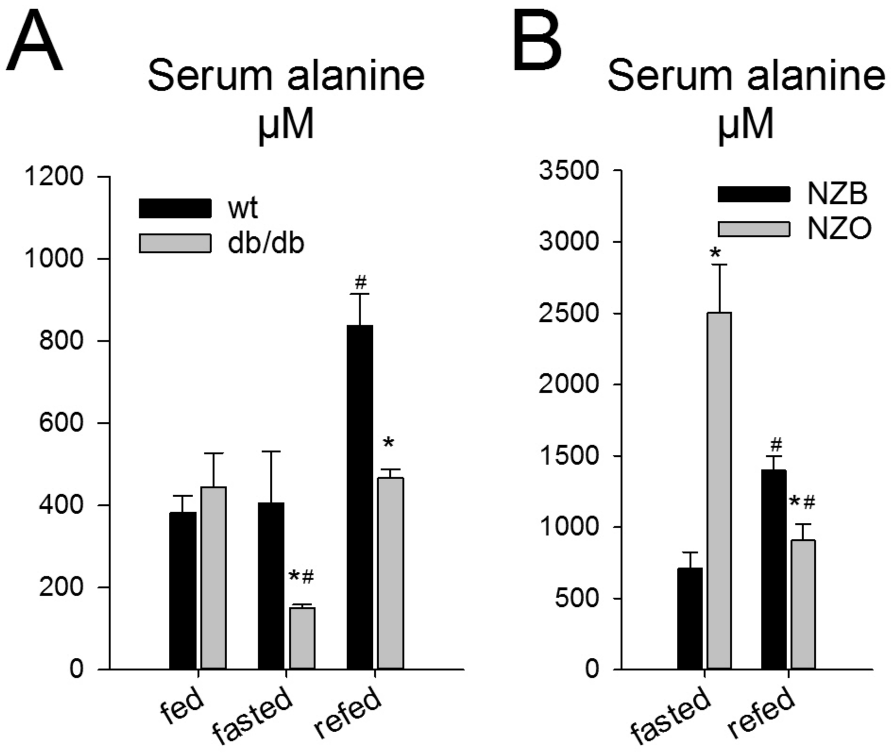


**Figure S1. Related to Figure 1.**

A: Serum alanine levels in wildtype C57Bl/6J (wt) and obese/diabetic BKS-db/db mice from ad libitum feeding (fed), 24h fasted (fasted) and 24h fasted, 6h refeeding (refed). Data are mean ± SEM, N=4/group. Effect of genotype: * $P > 0.05$. Effect of nutritional state vs. fed: # $P < 0.05$.

B: Serum alanine levels in New Zealand Black (NZB) and New Zealand Obese (NZO) mice from 24h fasting (fasted) and 24h fasted, 6h refeeding (refed). Data are mean ± SEM, N=4/group. Effect of genotype: * $P > 0.05$. Effect of nutritional state vs. fasted: # $P < 0.05$.

Statistical tests: A, B: 2-way ANOVA with Holm-Sidak posthoc hoc tests.

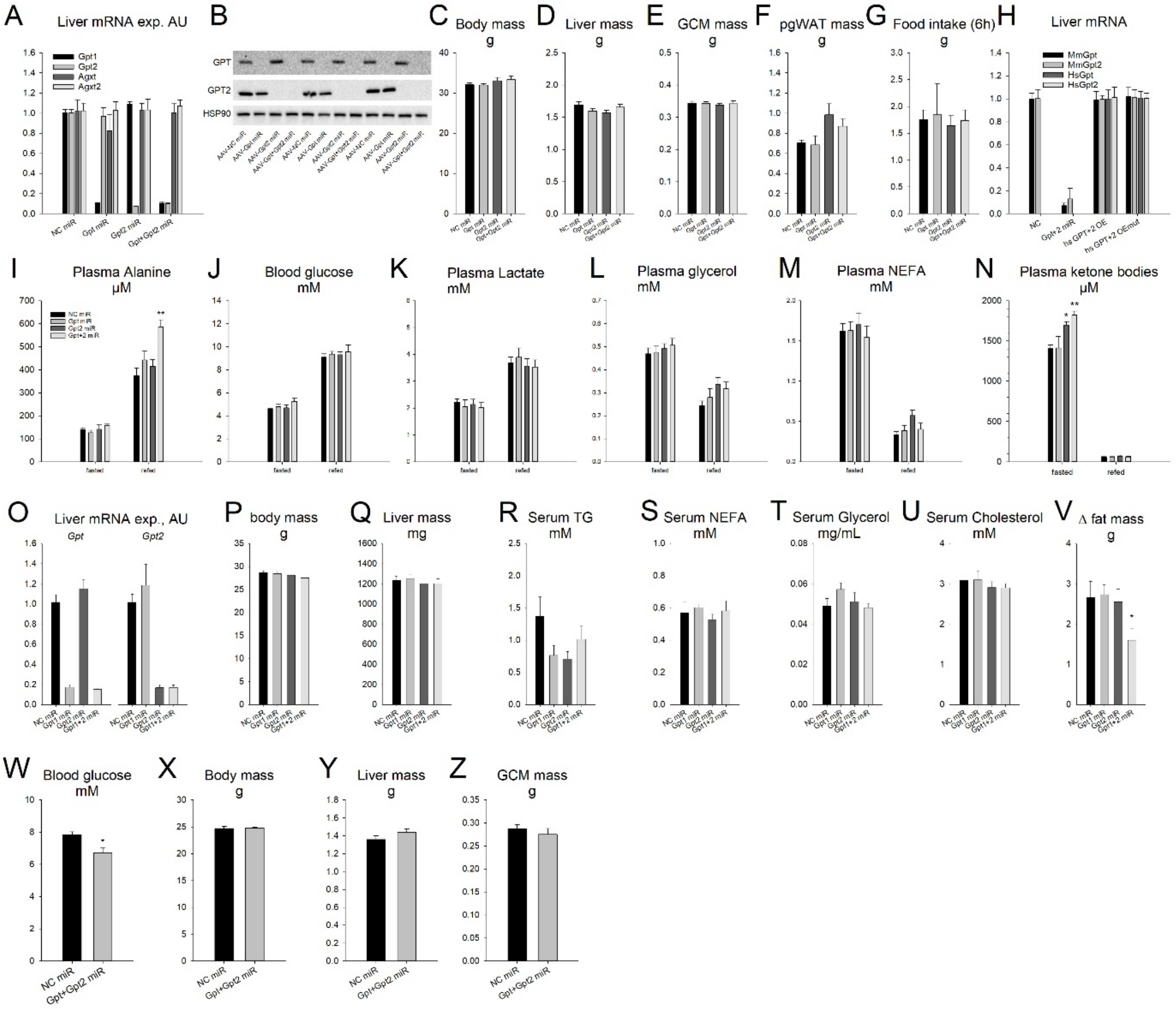


**Figure S2. Related to Figure 2.**

A: Liver mRNA expression from mice with hepatocyte selective AAV-miR mediated silencing of glutamic-pyruvic transaminase (Gpt) and alanine-glyoxylate aminotransferase (Agxt) isoforms. NC: negative control. miR: micro-RNA. Data are mean ± SEM, N = 6-7/group. Effect of miR vs. NC miR: * P < 0.05, ** P < 0.01, *** P < 0.001.

B: Representative western blots of GPT isoforms as well as the housekeeping protein heat shock protein 90 (HSP90) from mice as in A.

C: Body mass of mice as in A.

D: Liver mass of mice as in A.

E: Gastrocnemius complex skeletal muscle (GCM) mass of mice as in A.

F: Perigonadal white adipose tissue (pgWAT) mass of mice as in A.

G: Food intake during a 6h refeeding period following 24h fasting from mice as in A.

H: Liver mRNA levels from mice pre-treated with adeno-associated viruses to express a negative control micro-RNA (NC-miR) and green fluorescent protein (GFP), Gpt and GPT2-specfific miRs, human Gpt and Gpt2 mRNAs (HsGpt OE, HsGpt2 OE), and mutants of human Gopt and Gpt2 mRNAs to produce enzymatically inactive proteins (HsGptmut OE, HsGPT2mut OE). . Data are mean ± SEM, N = 6/ group. Different than NC: $^{a}$ $P < 0.05$, $^{aa}$ $P < 0.01$, $^{aaa}$ $P < 0.001$; different than Gpt+2 miR: $^{b}$ $P < 0.05$, $^{bb}$ $P < 0.01$, $^{bbb}$ $P < 0.001$. Different than hs GPT+2 OE: $^{c}$ $P < 0.05$, $^{cc}$ $P < 0.01$, $^{ccc}$ $P < 0.001$.

I: Plasma alanine levels from mice as in G.

J: Blood glucose levels from mice as in G.

K: Plasma lactate levels from mice as in G.

L: Plasma glycerol levels rom mice as in G.

M: Plasma non-esterified fatty acid (NEFA) levels from mice as in G.

N: Plasma ketone body levels from mice as in G.

O: Liver mRNA expression of glutamic-pyruvic transaminase (Gpt) isoforms in mice fed a ketogenic diet with hepatocyte selective AAV-miR mediated silencing of Gpt isoforms. NC: negative control. miR: micro-RNA. Data are mean ± SEM, N = 6/group. Effect of miR vs. NC miR: * $P < 0.05$, ** $P < 0.01$, *** $P < 0.001$.

P: Body mass from mice as in L.

Q: Liver mass from mice as in L.

R: Serum triglyceride (TG) levels from mice as in L.

S: Serum NEFA levels from mice as in L.

T: Serum glycerol levels from mice as in L.

U: Serum cholesterol levels from mice as in L.

V: The change (Δ) in fat mass as determined by ECHO-MRI before and after ketogenic diet feeding from mice as in L.

W: Blood glucose levels in mice in the ad libitum fed state following adaptation to a protein-enriched (80%E) diet with hepatocyte selective AAV-miR mediated silencing of glutamic-pyruvic transaminase

(Gpt) isoforms. NC: negative control. miR: micro-RNA. Data are mean ± SEM, N = 6/group. Effect of miR vs. NC miR: * P < 0.05, ** P < 0.01, *** P < 0.001.

X: Body mass from mice as in T.

Y: Liver mass from mice as in T.

Z: Gastrocnemius complex skeletal muscle (GCM) mass from mice as in T.

Statistical tests: C, D, E, F, P, Q, R, S, T, U, V: 1-way ANOVA with Holm-Sidak posthoc hoc tests. I, J, K, L, M, N: 2-way repeated measures ANOVA with Holm-Sidak posthoc hoc tests. W, X, Y, Z: Students t-tests.

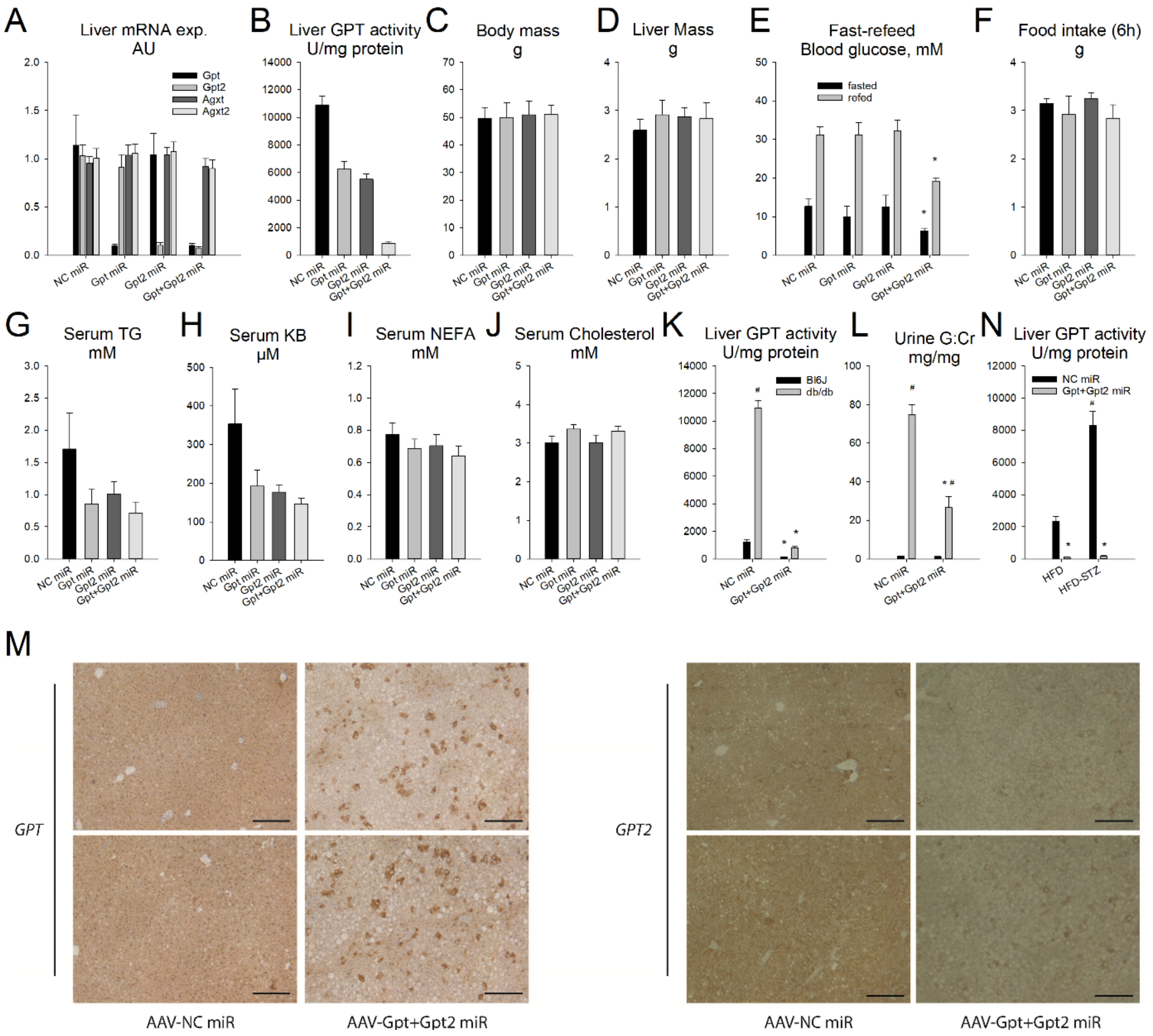


**Figure S3**: Related to Figure 3.

A: Liver glutamic-pyruvic transaminase isoform mRNA expression in obese/diabetic BKS-db/db mice with hepatocyte selective AAV-miR mediated silencing of Gpt isoforms. NC: negative control. miR: micro-RNA. Data are mean ± SEM, N = 6/group. Effect of miR vs. NC miR: * P < 0.05, ** P < 0.01, *** P < 0.001.

B: Liver GPT activity from mice as in A.

C: Body mass of mice as in A.

D: Liver mass of mice as in A.

E: Blood glucose levels after an overnight fast and following a 6h refeed from mice as in A. Food intake during a 6h refeeding period following 24h fasting from mice as in A.

F: Food intake during a 6h refeeding period following fasting from mice as in H.

G: Ad libitum fed serum triglyceride (TG) levels from mice as in A.

H: Serum ketone body (KB) levels from mice as in G.

I: Serum non-esterified fatty acid (NEFA) levels from mice as in G.

J: Serum cholesterol levels from mice as in G.

K: Liver GPT activity in lean C57Bl/6J (Bl6) and age-matched obese/diabetic BKS-db/db mice with hepatocyte selective AAV-miR mediated silencing of glutamic-pyruvic transaminase (Gpt) isoforms. NC: negative control. miR: micro-RNA. Data are mean ± SEM, N = 4/group. Effect of miR vs. NC miR: * $P < 0.05$, ** $P < 0.01$, *** $P < 0.001$. Effect of genotype: # $P < 0.05$, ## $P < 0.01$, ### $P < 0.001$.

L: Urinary glucose (G) to creatinine (Cr) ratio from mice as in K.

M: Liver glutamic-pyruvic transaminase (GPT) immunohistochemical staining of sections taken from BKS-db/db (db/db) mice with hepatocyte selective AAV-miR mediated silencing of glutamic-pyruvic transaminase (Gpt) isoforms from mice as in A. Shown are 2 representative images taken from 2 individual mice per group. Scale bar: 200µm.

N: Liver GPT activity of mice on an obesogenic high-fat diet with (HFD-STZ) or without (HFD) streptozocin (STZ) pre-treatment to exacerbate the progression of frank diabetes; with hepatocyte selective AAV-miR mediated silencing of glutamic-pyruvic transaminase (Gpt) isoforms. NC: negative control. Data are mean ± SEM, N = 6/group. Effect of miR vs. NC miR: * $P < 0.05$, ** $P < 0.01$, *** $P < 0.001$. Effect of STZ: # $P < 0.05$, ## $P < 0.01$, ### $P < 0.001$.

Statistical tests: C, D, F, G, H, I, J: 1-way ANOVA with Holm-Sidak posthoc hoc tests. E: 2-way repeated measures ANOVA with Holm-Sidak posthoc hoc tests. K, L, N: 2-way ANOVA with Holm-Sidak posthoc hoc tests.

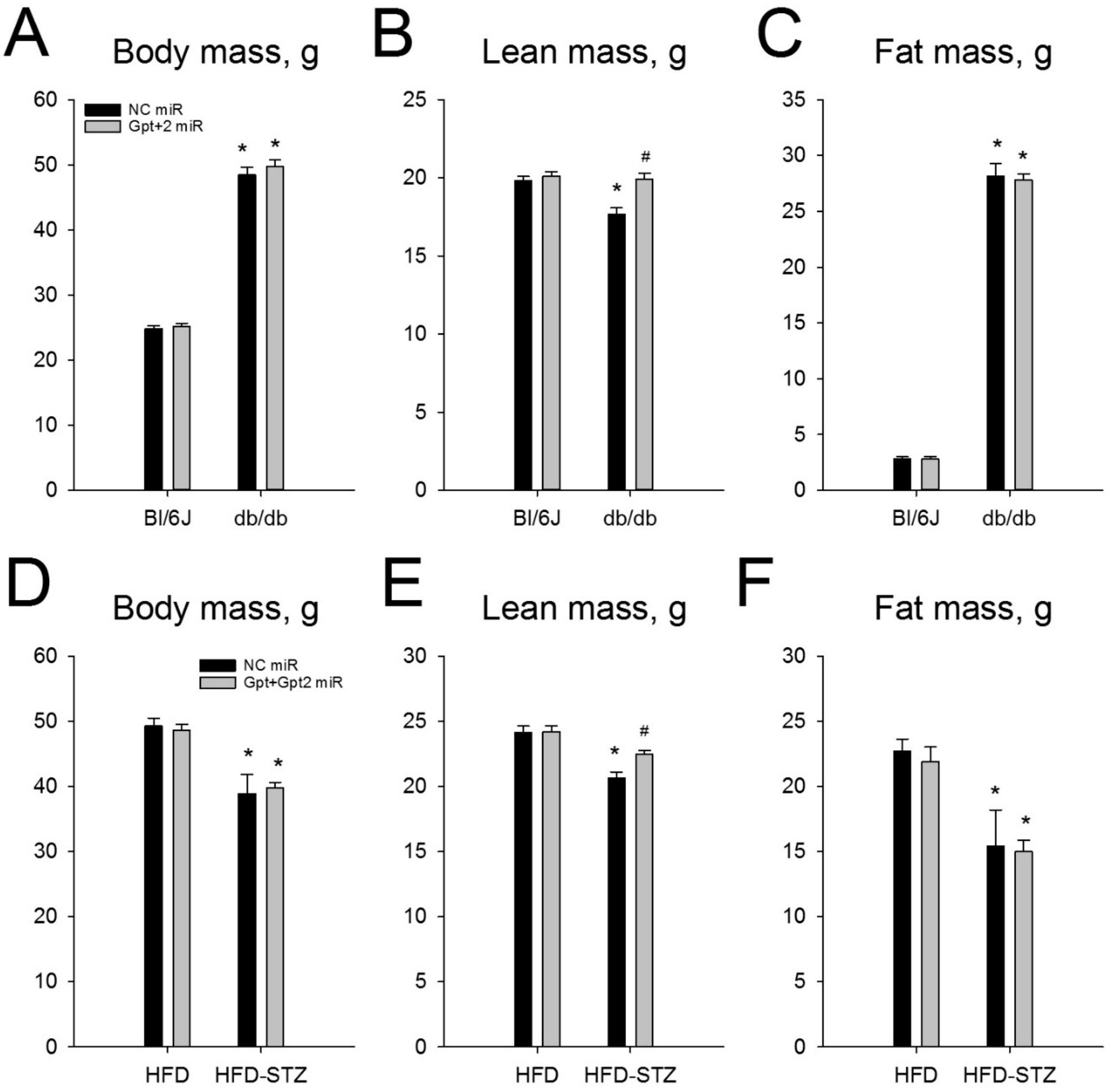


**Figure S4**. Related to Figure 4.

A: Body mass of lean C57Bl/6J (Bl6) and age-matched obese/diabetic BKS-db/db mice with hepatocyte selective AAV-miR mediated silencing of glutamic-pyruvic transaminase (Gpt) isoforms. NC: negative control. miR: micro-RNA. Data are mean ± SEM, N = 4/group. Effect of miR vs. NC miR: * P < 0.05, ** P < 0.01, *** P < 0.001. Effect of genotype: # P < 0.05, ## P < 0.01, ### P < 0.001.

B: Lean body mass as determined by ECHO-MRI of mice as in A.

C: Fat body mass as determined by ECHO-MRI of mice as in A.

D: Body mass of mice on an obesogenic high-fat diet with (HFD-STZ) or without (HFD) streptozocin (STZ) pre-treatment to exacerbate the progression of frank diabetes; with hepatocyte selective AAV-miR mediated silencing of glutamic-pyruvic transaminase (Gpt) isoforms. NC: negative control. Data are mean ± SEM, N = 6/group. Effect of miR vs. NC miR: * $P < 0.05$, ** $P < 0.01$, *** $P < 0.001$. Effect of STZ: # $P < 0.05$, ## $P < 0.01$, ### $P < 0.001$.

E: Lean body mass as determined by ECHO-MRI of mice as in D.

F: Fat body mass as determined by ECHO-MRI of mice as in D.

Statistical tests: A, B, C, D, E, F: 2-way ANOVA with Holm-Sidak posthoc hoc tests.

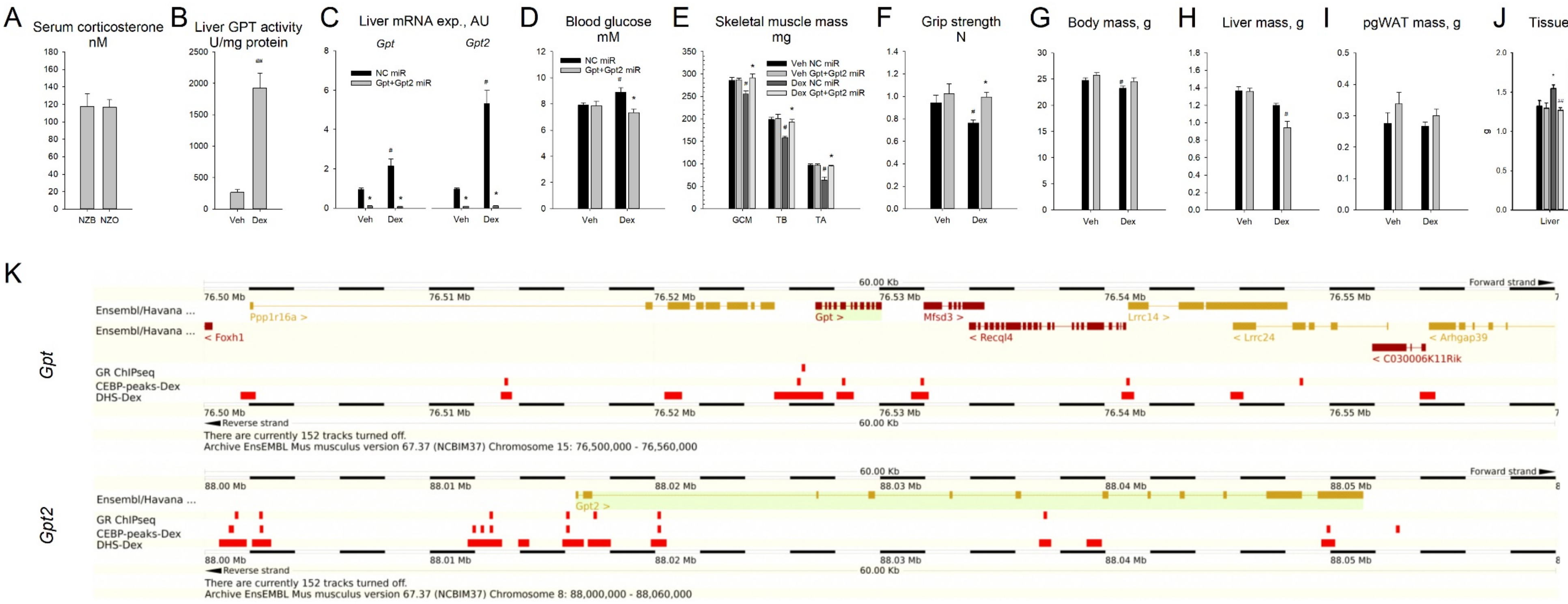

A
Serum corticosterone nM
NZB NZO
B
Liver GPT activity U/mg protein
Veh Dex
C
Liver mRNA exp., AU
Gpt
Gpt2
NC miR
Gpt+Gpt2 miR
Veh Dex Veh Dex
D
Blood glucose mM
NC miR
Gpt+Gpt2 miR
Veh Dex
E
Skeletal muscle mass mg
Veh NC miR
Veh Gpt+Gpt2 miR
Dex NC miR
Dex Gpt+Gpt2 miR
GCM TB TA
F
Grip strength N
Veh Dex
G
Body mass, g
Veh Dex
H
Liver mass, g
Veh Dex
I
pgWAT mass, g
Veh Dex
J
Tissue mass'
VEH NC-miR
DEX NC-miR
VEH Gpt+2-miR
DEX Gpt+2 miR
g
Liver pgWAT
K
Gpt
60.00 Kb
Forward strand
76.50 Mb 76.51 Mb 76.52 Mb 76.53 Mb 76.54 Mb 76.55 Mb
Ensembl/Havana ...
Ppp1r16a >
Gpt >
Mfsd3 >
Lrrc14 >
Ensembl/Havana ...
< Foxh1
< Recql4
< Lrrc24
< Arhgap39
< C030006K11Rik
GR ChIPseq
CEBP-peaks-Dex
DHS-Dex
Reverse strand
There are currently 152 tracks turned off.
Archive EnsEMBL Mus musculus version 67.37 (NCBIM37) Chromosome 15: 76,500,000 - 76,560,000
Gpt2
60.00 Kb
Forward strand
88.00 Mb 88.01 Mb 88.02 Mb 88.03 Mb 88.04 Mb 88.05 Mb
Ensembl/Havana ...
Gpt2 >
GR ChIPseq
CEBP-peaks-Dex
DHS-Dex
Reverse strand
There are currently 152 tracks turned off.
Archive EnsEMBL Mus musculus version 67.37 (NCBIM37) Chromosome 8: 88,000,000 - 88,060,000

**Figure S5**: Related to Figure 5.

A: Serum corticosterone levels in lean New Zealand Black (NZB) and obese/diabetic New Zealand Obese (NZO) mice. Data are mean ± SEM, N = 6/group.

D: Liver GPT activity of mice chronically treated with the synthetic glucocorticoid dexamethasone (Dex; 1mg/kg per day, 14d) or vehicle control (Veh). Data are mean ± SEM, N = 7/group. Effect of Dex: # P < 0.05, ## P < 0.01, ### P < 0.001.

C: Liver glutamic-pyruvic transaminase (Gpt) isoform mRNA expression of C57Bl/6N in mice chronically treated with the synthetic glucocorticoid dexamethasone (Dex; 1mg/kg per day, 14d) or vehicle control (Veh) with hepatocyte selective AAV-miR mediated silencing of glutamic-pyruvic transaminase (Gpt) isoforms. NC: negative control. miR: micro-RNA.. Data are mean ± SEM, N = 8/group. Effect of miR vs. NC miR: * P < 0.05, ** P < 0.01, *** P < 0.001. Effect of Dex: # P < 0.05, ## P < 0.01, ### P < 0.001.

D: Blood glucose levels of mice as in C.

E: Gastrocnemius complex (GCM), triceps brachii (TB) and tibialis anterior (TA) skeletal muscle mass' of mice as in C.

F: Forelimb grip strength of mice as in C.

G: Body mass of mice as in C.

H: Liver mass of mice as in C.

I: Perigonadal white adipose tissue (pgWAT) mass of mice as in C.

J: Liver and perigonal white adipose tissue mass' of mice chronically treated with the synthetic glucocorticoid dexamethasone (Dex; 1mg/kg per day, 14d) or vehicle control (Veh) with hepatocyte selective AAV-miR mediated silencing of glutamic-pyruvic transaminase (Gpt) isoforms. NC: negative control. miR: micro-RNA. Data are mean ± SEM, N = 8/group. Effect of miR vs. NC miR: * P < 0.05, ** P < 0.01, *** P < 0.001. Effect of Dex: # P < 0.05, ## P < 0.01, ### P < 0.001.

K: ENSEMBL genome browser images of DNAse hypersensitivity sites as well as CEBPb and GR ChIP-seq peaks of the Gpt and Gpt2 gene, and flanking regions, in mouse liver.

Statistical tests: A, B: Students t-tests. C, D, E, F, G, H, I, J: 2-way ANOVA with Holm-Sidak posthoc hoc tests.

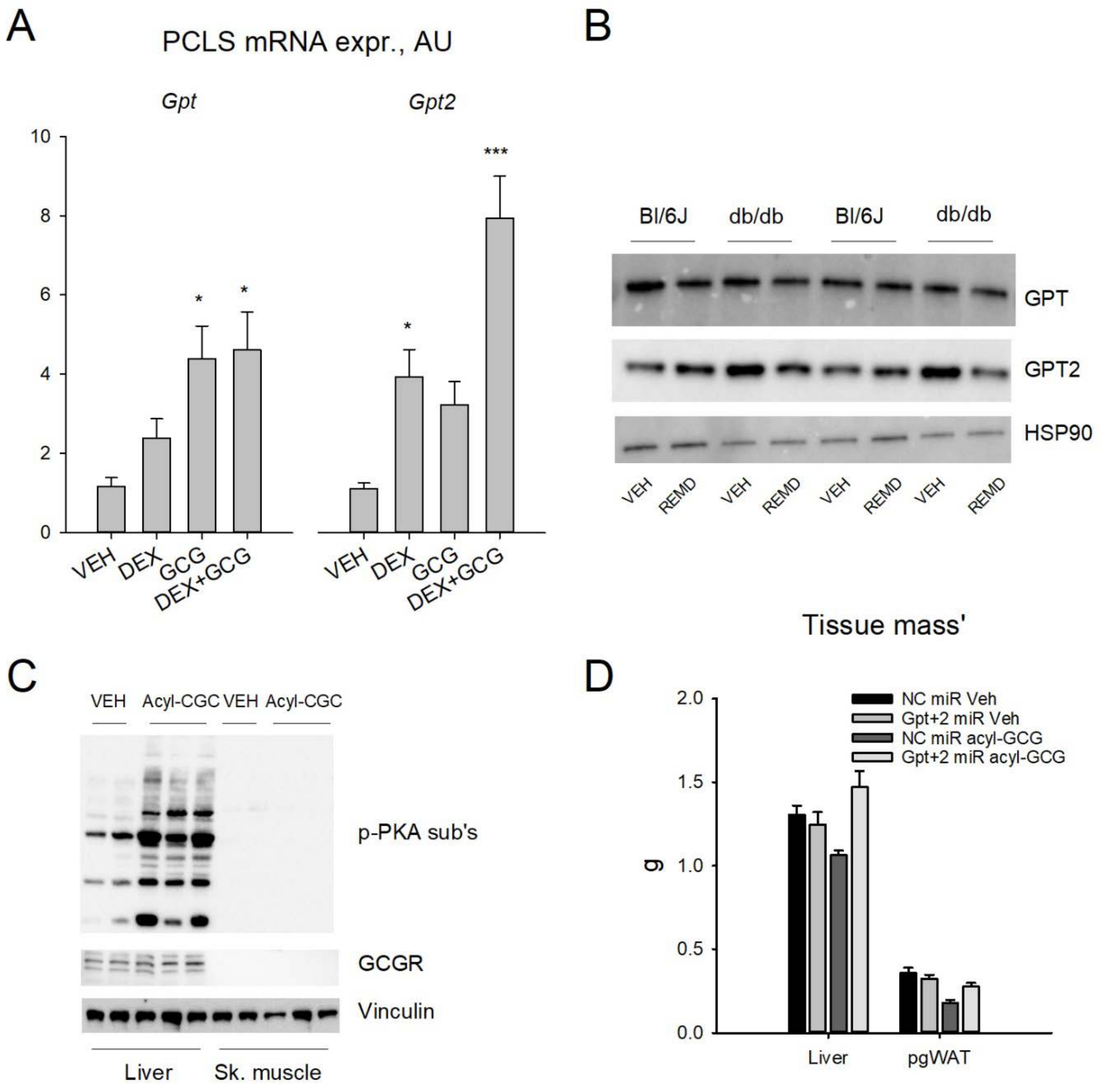


**Figure S6**: Related to Figure 6.

A: Messenger RNA (mRNA) levels of glutamic-pyruvic transaminase (Gpt) and Gpt2 in precision cut liver slices treated ex vivo with the synthetic glucocorticoid dexamethasone (DEX), glucagon (GCG) or a combination thereof (DEX+GCG). N= 6 individual slices per treatment group. One way ANOVA: different than VEH: * $p < 0.05$, ** $p < 0.01$, *** $p < 0.001$.

B: Western blot images of liver glutamic-pyruvic transaminase (GPT), GPT2, and loading control heat shock protein 90 (HSP90) of C57Bl6/J (Bl6/J) and obese/diabetic C57BKS mice with homozygous leptin receptor mutation (BKS-db/db) chronically treated with a glucagon receptor antagonist (REMD).

C: Western blot images of liver and skeletal muscle phospho-protein kinase A motif protein substrates (p-PKA sub's), glucagon receptor (GCGR), and loading control Vinculin from mice acutely treated with acyl-glucagon (Acyl-GCG) or Vehicle (VEH).

D: Liver and perigonadal white adipose tissue (PGWAT) mass' of mice of C57Bl/6J mice chronically treated with the acyl-glucagon (acyl-GCG; 1 nmol/g/d) or vehicle control (Veh) with hepatocyte selective AAV-miR mediated silencing of glutamic-pyruvic transaminase (Gpt) isoforms. NC: negative control. miR: micro-RNA. Data are mean ± SEM, N = 8/group. Effect of miR vs. NC miR: * $P < 0.05$, ** $P < 0.01$, *** $P < 0.001$. Effect of Dex: # $P < 0.05$, ## $P < 0.01$, ### $P < 0.001$.

Statistical tests: A, D: 1-way ANOVA with Holm-Sidak posthoc hoc tests.

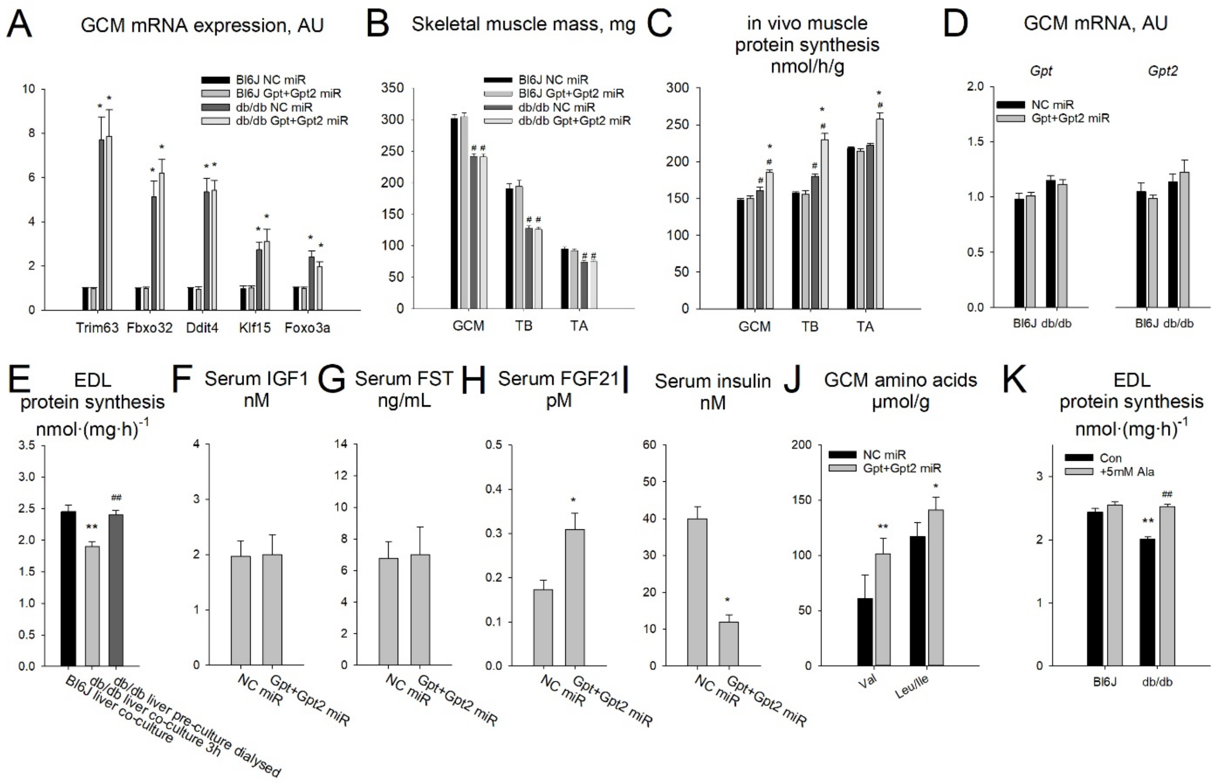


**Figure S7**: Related to Figure 7.

A: Gastroconemius complex skeletal muscle mRNA expression of transcripts of atrophy related genes Trim63 (aka Murf1), Fbxo32 (aka MAFbx or Atrogin-1), Ddit4 (aka Redd1), Klf15 and Foxo3a from lean C57Bl/6J (Bl6) and age-matched obese/diabetic BKS-db/db mice with hepatocyte selective AAV-miR mediated silencing of glutamic-pyruvic transaminase (Gpt) isoforms. Data are from mice 6 wk after AAV administration. NC: negative control. miR: micro-RNA. GCM: gastrocnemius complex muscle. TB: Triceps brachii. TA: tibialis anterior. Data are mean ± SEM, N = 4/group. Effect of miR vs. NC miR: * $P < 0.05$, ** $P < 0.01$, *** $P < 0.001$. Effect of genotype: # $P < 0.05$, ## $P < 0.01$, ### $P < 0.001$.

B: Skeletal muscle mass' of overnight fasted, 4h refed lean C57Bl/6J (Bl6) and age-matched obese/diabetic BKS-db/db mice with hepatocyte selective AAV-miR mediated silencing of glutamic-pyruvic transaminase (Gpt) isoforms. Study was conducted 10d after AAV administrations. NC: negative control. miR: micro-RNA. GCM: gastrocnemius complex muscle. TB: Triceps brachii. TA: tibialis anterior. Data are mean ± SEM, N = 4/group. Effect of miR vs. NC miR: * $P < 0.05$, ** $P < 0.01$, *** $P < 0.001$. Effect of genotype: # $P < 0.05$, ## $P < 0.01$, ### $P < 0.001$.

C: In vivo protein synthesis rate calculated from mixed muscle $^3$H-phenylalanine incorporation of mice as in B.

D: Gastrocnemius complex skeletal muscle (GCM) Gpt isoform mRNA expression in lean C57Bl/6J (Bl6) and age-matched obese/diabetic BKS-db/db mice with hepatocyte selective AAV-miR mediated silencing of glutamic-pyruvic transaminase (Gpt) isoforms. NC: negative control. miR: micro-RNA. Data are mean ± SEM, N = 4/group. Effect of miR vs. NC miR: * $P < 0.05$, ** $P < 0.01$, *** $P < 0.001$. Effect of genotype: # $P < 0.05$, ## $P < 0.01$, ### $P < 0.001$.

E: Ex vivo extensor digitorum longus (EDL) skeletal muscle protein synthesis rate during co-culture and cross-co-culture with liver slices. Tissues were taken from lean C57Bl/6J (Bl6) and age-matched obese/diabetic BKS-db/db mice with hepatocyte selective AAV-miR mediated silencing of glutamic-pyruvic transaminase (Gpt) isoforms. In one condition, media was collected after culture with db/db liver for 3h after which it was dialyzed (1 KDa cutoff) against normal media to normalize metabolites such as amino acids but retain large molecules such as peptides. NC: negative control. miR: micro-RNA. Data are mean ± SEM, N = 3/group with 2 technical replicates per treatment condition. Effect of Liver miR: * $P < 0.05$, ** $P < 0.01$, *** $P < 0.001$. Effect of muscle genotype: # $P < 0.05$, ## $P < 0.01$, ### $P < 0.001$.

F: Serum insulin like growth factor 1 (IGF1) levels of BKS-db/db mice as in D.

G: Serum follistatin (FST) levels of BKS-db/db mice as in D.

H: Serum fibroblast growth factor 21 (FGF21) levels of BKS-db/db mice as in D.

I: Serum insulin levels of BKS-db/db mice as in D.

J: Gastrocnemius complex skeletal muscle (GCM) Valine (Val) and Leucine/Isoleucine (Leu/Ile) concentrations in obese/diabetic BKS-db/db mice with hepatocyte selective AAV-miR mediated silencing of glutamic-pyruvic transaminase (Gpt) isoforms. NC: negative control. miR: micro-RNA. Data are mean ± SEM, N = 6/group. Effect of miR vs. NC miR: * $P < 0.05$, ** $P < 0.01$, *** $P < 0.001$.

K: Ex vivo extensor digitorum longus (EDL) skeletal muscle protein synthesis rate during co-culture and cross-co-culture with liver slices with (+5mM Ala) or without (Con) treatment with 5mM alanine in the media. Tissues were taken from lean C57Bl/6J (Bl6) and age-matched obese/diabetic BKS-db/db mice. NC: negative control. miR: micro-RNA. Data are mean ± SEM, N = 3/group with 2 technical replicates per treatment condition. Effect of muscle genotype: * $P < 0.05$, ** $P < 0.01$, *** $P < 0.001$. Effect of Co8b: # $P < 0.05$, ## $P < 0.01$, ### $P < 0.001$.

Statistical tests: A, B, C, D, J: 2-way ANOVA with Holm-Sidak posthoc hoc tests. E, F, G, H, K: Student's t-test. I: 1-way ANOVA with Holm-Sidak posthoc hoc tests.
